# A Novel Approach-Avoidance Task to Study Decision Making Under Outcome Uncertainty

**DOI:** 10.1101/2025.07.12.663075

**Authors:** Ziwei Cheng, Nadja R. Ging-Jehli, Maisy Tarlow, Joonhwa Kim, Henry W. Chase, Manan Arora, Lisa Bonar, Ricki Stiffler, Alex Grattery, Simona Graur, Michael J. Frank, Mary L. Phillips, Amitai Shenhav

## Abstract

To behave adaptively, people need to integrate information about probabilistic outcomes and balance drives to approach positive outcomes and avoid negative outcomes. However, questions remain about how uncertainty in positive and negative outcomes influence approach-avoid decision-making dynamics. To fill this gap, we developed a novel Probabilistic Approach Avoidance Task (PAAT) and characterized behavior in this task using sequential sampling models In this task, participants (Study 1: blinded mixed clinical sample N=34; Study 2: online nonpsychiatric sample N = 58) made a series of choices between pairs of options, each consisting of variable probabilities of reaching a positive outcome (monetary reward) and of reaching a negative outcome (aversive image). Participants tended to choose options that maximized the likelihood of reward and minimized the likelihood of aversive outcomes. Moreover, the weights they placed on each of these differed for choices where these likelihoods were in opposition (i.e., the riskier option was also more rewarding; incongruent trials) relative to when these were aligned (congruent trials). Computational modeling revealed that the relative influence of rewarding and aversive outcomes on choice was captured by differences in the rate of decision-relevant information accumulation. These modeling results were validated with a series of model comparisons and posterior predictive checks, demonstrating that our sequential sampling models reliably captured our behavioral data. Together, these findings improve our understanding of the influence of motivational conflict, outcome type, and levels of uncertainty on approach-avoid decision-making.

---

Everyday behavior is guided both by the drive to seek desirable outcomes (approach motivation) and the drive to prevent undesirable outcomes (avoidance motivation) (Atkinson, 1957; Schneirla, 1959; Elliot, 2013). Oftentimes, these motivations compete with one another – to increase their chances of achieving a positive outcome, a person must risk higher likelihoods of an aversive outcome occurring. In such cases of *approach-avoid conflict* (AAC), the person must simultaneously weigh the valence, magnitude, and likelihood of expected outcomes to make a decision (Kirlic et al., 2017; Ging-Jehli et al., 2024). Suboptimal evaluation or integration of these factors may lead to maladaptive motivated behavior and prolonged negative mood (Loijen et al., 2020; Trew, 2011; Angelakis & Pseftogianni, 2021; Akbari et al., 2022).

Thus, understanding the cognitive mechanisms underpinning approach-avoid decision-making is of great theoretical and translational importance. Previous studies have sought to fill this gap with a variety of tasks in which participants face trade-offs between positive and negative outcomes, primarily in cases where those outcomes are expected to occur with certainty if the relevant action is selected (e.g., Aupperle et al., 2011; Ironside et al., 2020; Klaassen et al., 2021; Talmi et al., 2009; Zorowitz et al., 2019; Ging-Jehli et al., 2024; Table 1). Real-world approach-avoid conflicts, by contrast, typically involve outcomes that are uncertain (Monosov, 2020). Some work has begun to address this by having outcomes (e.g., shock versus rewards) occur with a fixed probability, and varying the magnitude of those outcomes (Ging-Jehli et al., 2024; Klaassen et al., 2021), or by varying the uncertainty and magnitude of approach-related outcomes, but not avoidance-related outcomes (e.g., Sierra-Mercado et al., 2015; Ironside et al., 2020; Bach 2015). However, this still leaves open the question of how approach-avoidance decisions vary as a function of outcome likelihood, that is, as rewarding and aversive outcomes vary from having a high likelihood of occurring to having a high likelihood of not occurring.

**Table 1.**
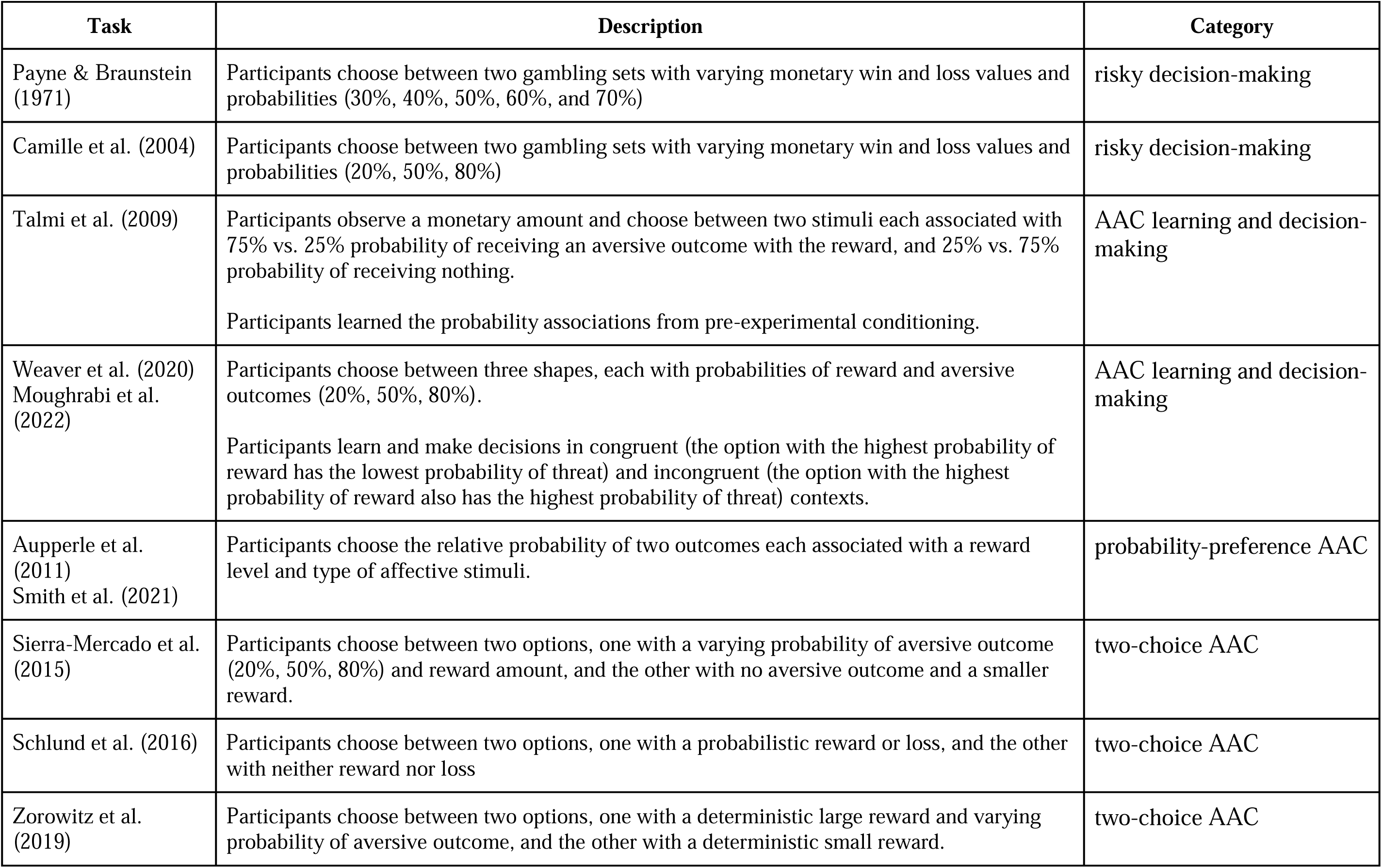
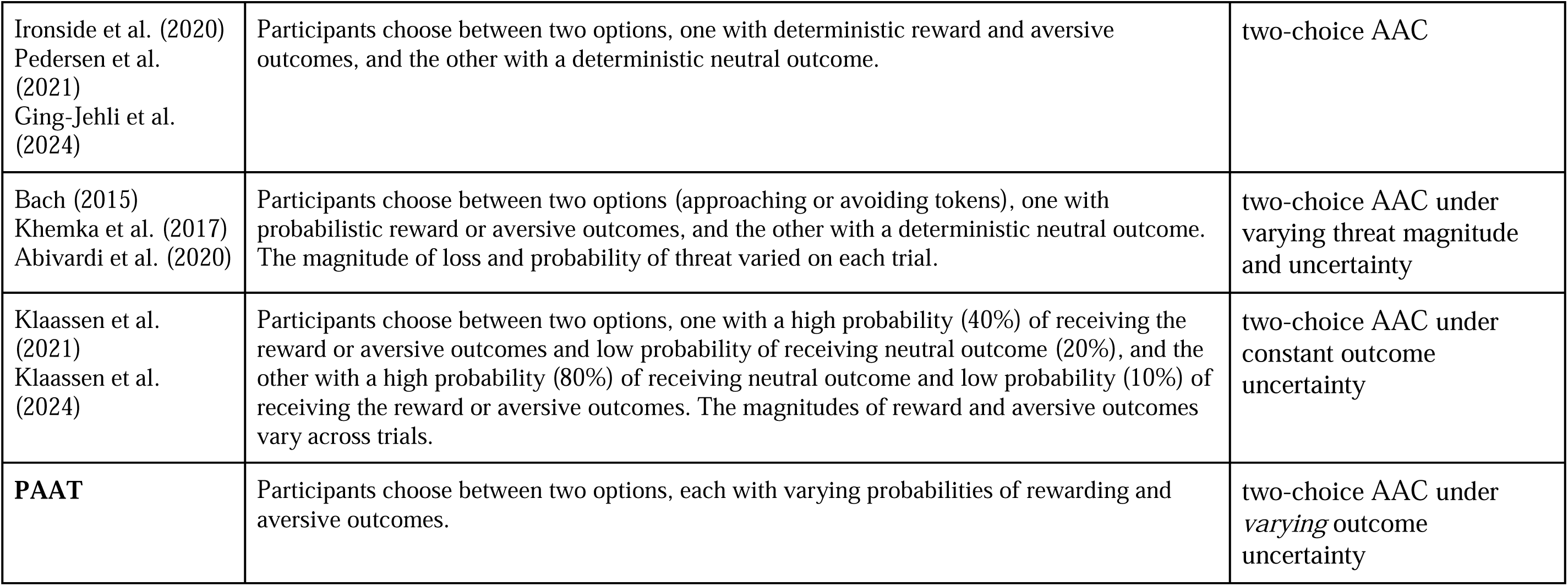
Descriptions of PAAT and previous relevant behavioral paradigms that investigated Approach-Avoid Conflict.

PROBABILISTIC APPROACH AVOIDANCE TASK
| Task | Description | Category |
| --- | --- | --- |
| Payne & Braunstein (1971) | Participants choose between two gambling sets with varying monetary win and loss values and probabilities (30%, 40%, 50%, 60%, and 70%) | risky decision-making |
| Camille et al. (2004) | Participants choose between two gambling sets with varying monetary win and loss values and probabilities (20%, 50%, 80%) | risky decision-making |
| Talmi et al. (2009) | Participants observe a monetary amount and choose between two stimuli each associated with 75% vs. 25% probability of receiving an aversive outcome with the reward, and 25% vs. 75% probability of receiving nothing.<br>Participants learned the probability associations from pre-experimental conditioning. | AAC learning and decision-making |
| Weaver et al. (2020)<br>Moughrabi et al. (2022) | Participants choose between three shapes, each with probabilities of reward and aversive outcomes (20%, 50%, 80%).<br>Participants learn and make decisions in congruent (the option with the highest probability of reward has the lowest probability of threat) and incongruent (the option with the highest probability of reward also has the highest probability of threat) contexts. | AAC learning and decision-making |
| Aupperle et al. (2011)<br>Smith et al. (2021) | Participants choose the relative probability of two outcomes each associated with a reward level and type of affective stimuli. | probability-preference AAC |
| Sierra-Mercado et al. (2015) | Participants choose between two options, one with a varying probability of aversive outcome (20%, 50%, 80%) and reward amount, and the other with no aversive outcome and a smaller reward. | two-choice AAC |
| Schlund et al. (2016) | Participants choose between two options, one with a probabilistic reward or loss, and the other with neither reward nor loss | two-choice AAC |
| Zorowitz et al. (2019) | Participants choose between two options, one with a deterministic large reward and varying probability of aversive outcome, and the other with a deterministic small reward. | two-choice AAC |

|  |  |  |
| --- | --- | --- |
| Ironside et al. (2020)<br>Pedersen et al. (2021)<br>Ging-Jehli et al. (2024) | Participants choose between two options, one with deterministic reward and aversive outcomes, and the other with a deterministic neutral outcome. | two-choice AAC |
| Bach (2015)<br>Khemka et al. (2017)<br>Abivardi et al. (2020) | Participants choose between two options (approaching or avoiding tokens), one with probabilistic reward or aversive outcomes, and the other with a deterministic neutral outcome. The magnitude of loss and probability of threat varied on each trial. | two-choice AAC under varying threat magnitude and uncertainty |
| Klaassen et al. (2021)<br>Klaassen et al. (2024) | Participants choose between two options, one with a high probability (40%) of receiving the reward or aversive outcomes and low probability of receiving neutral outcome (20%), and the other with a high probability (80%) of receiving neutral outcome and low probability (10%) of receiving the reward or aversive outcomes. The magnitudes of reward and aversive outcomes vary across trials. | two-choice AAC under constant outcome uncertainty |
| **PAAT** | Participants choose between two options, each with varying probabilities of rewarding and aversive outcomes. | two-choice AAC under *varying* outcome uncertainty |

To address these gaps and obtain computational insights into the cognitive mechanisms of approach-avoid decision-making under outcome uncertainty, the current study developed a Probabilistic Approach-Avoid Task (PAAT; Figure 1). On each trial in this task, participants choose between 2 options, with each option carrying some likelihood of a positive outcome (monetary reward) and some likelihood of a negative outcome (aversive image). The probabilities of positive and negative outcomes varied independently. Across trials, the task also varied whether approach and avoidance motivations were in alignment (i.e., safer options were also more likely to be rewarded; *congruent* trials) or were in opposition (i.e., risky options were more likely to be rewarded; *incongruent* trials). This additional manipulation allowed us to examine the evaluation of reward and aversive outcome probability under different contexts, comprehensively characterizing choice dynamics across conflict domain and conflict levels (Talmi et al., 2009; Aupperle et al., 2011; Weaver et al., 2020).

**Figure 1.**
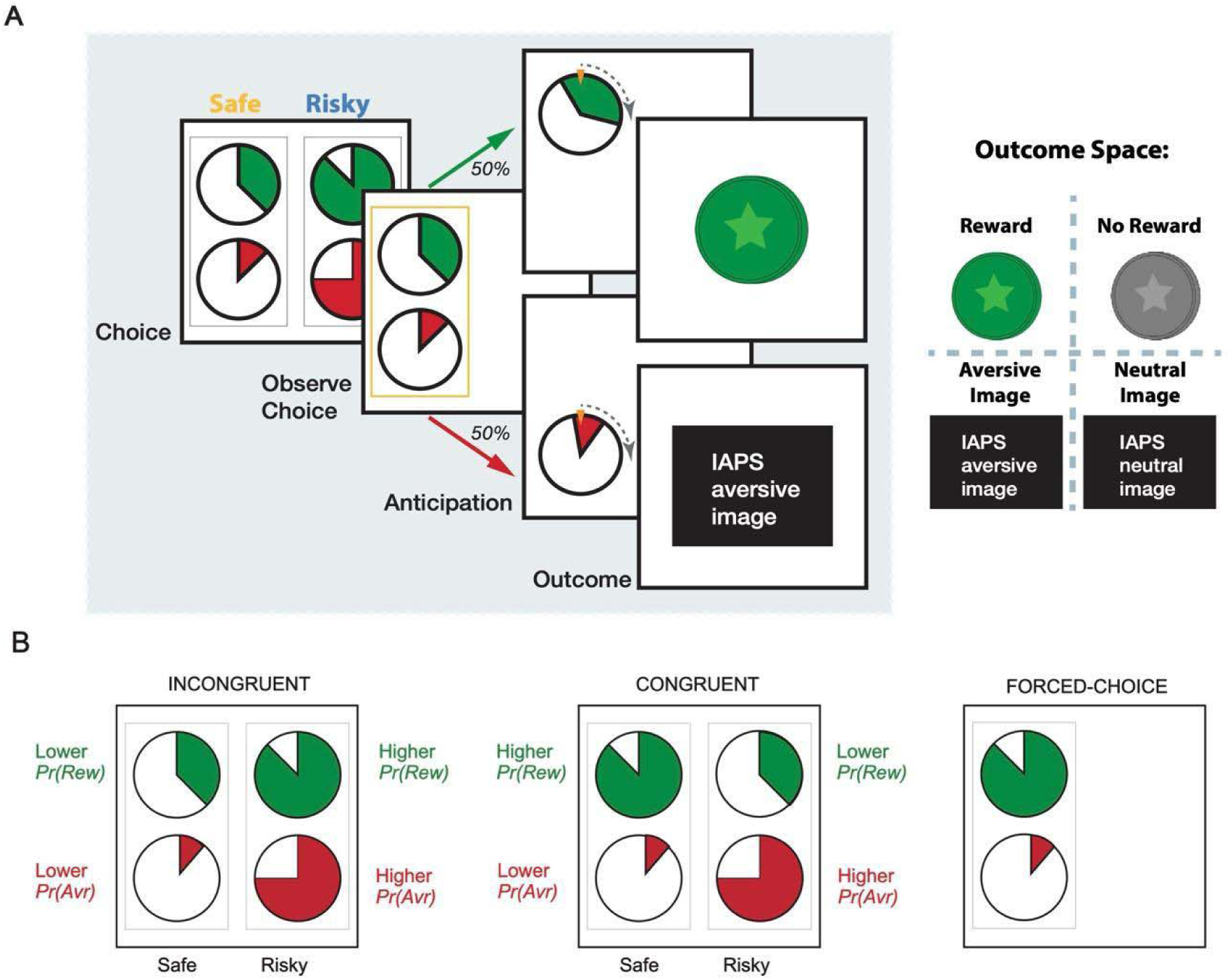
Probabilistic Approach Avoidance Task. (A) On each trial, each of 2 options had a probability of positive (money) and a negative outcome (aversive image). Participants chose which pair of wheels they prefer in the choice phase, one wheel was chosen at random to remain on the screen from the chosen pair, and participants viewed the outcome determined by the wheel spin. (B) In incongruent trials, the risky option had a higher probability of a rewarding outcome, whereas for congruent trials the risky option had a lower probability of a rewarding outcome. The task also included forced choice trials (24 total) where participants had only one option (a pair of a positive outcome probability and a negative outcome probability) and were cued to choose the option. These trials would be relevant for future neuroimaging analysis to understand outcome probability processing and were excluded from current behavioral analysis.

To characterize the processes and dynamics underlying these probabilistic approach-avoid decisions, we fit the data with sequential sampling models (SSMs). By jointly accounting for response times (RT) and response frequencies, SSMs provide an algorithmic account of how evidence in value-based decision accumulates during the decision process (e.g., choosing to approach vs. avoid) (Westbrook et al., 2020; Frank, 2025; Turner et al., 2018). These models were originally primarily applied to characterize perceptual decision-making processes and have since been shown to accurately capture behavior across a wide range of domains, including value-based decision-making tasks in neuroscience and economics (Gluth et al., 2012; Krajbich & Rangel, 2011; Milosavljevic et al., 2010; Krajbich, 2019; Ging-Jehli et al., 2023, 2025). Previous studies have also demonstrated within-trial accumulation of option values or incentives in neural recordings (Rich & Wallis, 2016; Doi et al., 2020), and evidence accumulation models that formalize these latent dynamics can capture more information than classic analyses or other distribution models that primarily focus on response times (Clithero, 2018; Forstmann et al., 2016; Fengler et al., 2021; Ging-Jehli et al., 2021). Moreover, they have been applied in clinical contexts, such as psychiatric diagnosis and assessment (e.g., Ging-Jehli et al., 2021, 2022; Wiecki et al., 2015; Pedersen et al., 2021). Through these applications, the modeling parameters have gained well-established psychological interpretations (Forstmann et al., 2016; Ging-Jehli et al., 2021), enabling us to formulate testable hypotheses about the influence of task variables (e.g., uncertainty, outcome type) on specific decision parameters. To effectively establish the underlying decision process and disentangle the influence of each task variable, we compared SSMs with different specifications about the decision dynamics (e.g., drift diffusion model, linear collapse model).

Based on past findings (Talmi et al., 2009; Klaassen et al., 2021; Ging-Jehli et al., 2024), we predicted that people would evaluate and integrate information about the motivational context, probability of reward, and probability of aversive outcome into decision-making. Across two studies (Study 1: blinded balanced sample of patients with obsessive-compulsive disorder and nonpsychiatric controls; Study 2: online nonpsychiatric sample), we found that decisions on incongruent trials were determined by both the probability of reward and the probability of aversive outcomes, whereas congruent trials were primarily influenced by the probability of reward.Our computational models were able to account for both choices and RT distributions in the task, and the best-fitting model revealed that the probability that a choice would lead to reward increased drift rates in favor of that choice, whereas the probability of a choice leading to an aversive outcome had the opposite effect. Collectively, these findings help to validate our new paradigm and provide insights into the relative contributions of outcome type and likelihood in approach-avoid decision making.

## Methods and Materials

### Participants

#### Study 1

Thirty-four adults were recruited from Pittsburgh, PA area and all participants provided informed consent to participate in the study (Gender: 71% females; Age: 25.2 ± 4.5 years; OCD diagnoses: 50%; Supplement Table S1). The sample was part of an ongoing study aimed at recruiting a balanced sample of patients with obsessive-compulsive disorder (OCD) and individuals without mental health conditions (see Supplement exclusion and inclusion criteria). The diagnostic status of the participants is blinded to the investigators, and clinical variables are not considered in the current investigation.

#### Study 2

Power analysis (Murayama et al., 2022) on Study 1 found that 43 participants were required to detect the smallest effect of interest (motivation congruency effect on response time; t = 2.57) with 80% power, and we collected 58 participants online through Prolific. 8 participants were excluded for choosing the option with lower reward and higher aversive probability on more than 20% of congruent trials or completing fewer than 80% of trials. The remaining 50 participants (Gender: 46% Female; Age: 37±10 years; Supplement Table S1) were included in all of our analyses.

### Task

We developed a novel Probabilistic Approach Avoidance Task (PAAT, Figure 1) to parametrically vary uncertainty in positive and negative outcomes respectively. On each two-alternative choice trials, participants chose between pairs of options, and each option included a probability of a rewarding outcome ($0.50 for Study 1 and $0.10 for Study 2 monetary gain indicated by a green token) and a probability of an aversive outcome (aversive image). Outcome probabilities were represented as wedges in a wheel, where green represented rewarding outcome probability and red represented aversive outcome probability. Each trial has 3 main phases: choice, anticipation and outcome. After participants chose an option in each trial (choice phase), one of the wheels (green or red) in the chosen pair would be randomly selected (50% probability) to remain on the screen (choice shown). A pointer appeared at the top of the selected wheel, which spun for 4 seconds (anticipation phase), and participants were shown the outcome determined by the wheel spin for 2 seconds (outcome phase). The possible outcomes of the green wheel included reward (if stopped at the green wedge) or no reward (if stopped at the empty wedge), and the possible outcomes of the red wheel included aversive image (if stopped at the red wedge) or neutral image (if stopped at the empty wedge). The response deadline during the choice period was 6 seconds in each trial, and participants would view an aversive image if they did not make a response in time. Across trials, we varied the probability of reward and aversive outcomes associated with each option (10-90% probability range at 20% increments) such that the options varied in the extent to which one was more likely to yield a reward and/or more likely to yield an aversive outcome. We counter-balanced the relative probabilities between the left and right options across trials. Trials in which one option had both a higher probability of reward and a higher probability of an aversive outcome were classified as *incongruent*, placing the approach-avoid motivations into conflict. In contrast, trials where one option had a higher probability of reward and a lower probability of an aversive outcome were classified as *congruent*.

### Procedures

#### Study 1

Participants completed a Probabilistic Approach and Avoidance Task (PAAT) as part of a larger ongoing clinical study. Before the task started, participants received detailed instructions and rated stimuli from the International Affective Picture System (Lang et al., 2008) for their valence and arousal (same images across participants; see Supplement for analysis of IAPS ratings). The present study was centered on the behavioral data from PAAT. Additional experimental investigations using overlapping samples and targeting independent hypotheses will be reported elsewhere. The task included a total of 120 trials, divided into 6 blocks of 20 trials each. Within each block, participants completed 16 two-alternative choice trials and 4 forced-choice trials (Figure 1). Due to a change in task version, 18 participants completed the task where the percentage of incongruent trials ranged between 42% and 63%, and 16 participants completed the task where the percentage was 69%. The average percentage of incongruent and congruent trials in the sample was 60% and 40%. Supplementary analyses (Table S2) showed that results (e.g., effects of trial-level probabilities on task behavior) did not differ with these ratios on congruent (ps > .06) or incongruent trials (ps > .07), and the main analyses collapsed data across all participants. Participants were informed that they would receive a monetary bonus based on rewards gained in the task. At the end of the study, all participants were debriefed and received a $25 bonus for completing the session.

#### Study 2

Participants completed PAAT online (implemented within the PsiTurk framework). Same as Study 1, participants received detailed instructions and rated IAPS stimuli for their valence and arousal. The task included a total of 96 (60 incongruent and 36 congruent) trials, divided into 6 blocks of 16 two-alternative choice trials each. At the end of the study, all participants were debriefed and received a monetary bonus based on rewards gained in the task.

### Behavioral Analysis

On each trial, we defined the risky option as the option with a higher probability of an aversive outcome (and safe option as the one with a lower probability of an aversive outcome). For incongruent trials, the risky option was the one that had a higher probability of a reward, whereas for congruent trials the safe option also entailed higher likelihood of reward.

We examined trial-level decision and response time in two ways: (1) using trial-wise probability of rewarding and aversive outcomes (*Pr(Rew)*_*risky*_, *Pr(Rew)*_*safe*_, *Pr(Avr)*_*risky*_, and *Pr(Avr)*_*safe*_) and (2) using the relative differences in probability of rewarding or aversive outcome between the risky and safe options (*Rel Rew = Pr(Rew)*_*risky*_*, - Pr(Rew)*_*safe*_; *Rel Avr = Pr(Avr)*_*risky*_ *- Pr(Avr)*_*safer*_). Model comparisons (in both regression and computational models) found that the relative values captured variability in behavior across PAAT trials with fewer parameters, and we focus on *Rel Rew* and *Rel Avr* in the main analysis. Detailed regression model comparisons and computational model comparisons, including alternative model specifications and results, are provided in the Supplementary Materials (Table S4 and S5).

Across *Rel Rew* (ranges from -0.8 to 0.8) and *Rel Avr* (ranges from 0.1 to 0.8), we presented variations in choice frequencies, average response time (RT), and RT quantiles across trials. In addition, we fitted linear mixed-effects regression models with the lme4 package (Bates et al., 2015) in R (R Core Team, 2023) to quantify the influence of outcome type and degree of outcome uncertainty on approach decisions (choosing the risky option) and RT.

We fitted the regression models separately on incongruent and congruent trials, as well as on all trials with congruency included as an additional predictor (binary-coded with incongruent trials as the reference group). To model trial-wise decision (choose risky = 1, choose safe = 0), we included *Rel Rew* and *Rel Avr* as predictors. To model trial-level (log-transformed) RT, we included *Rel Avr* and trial-level decision (choose risky =1, choose safe=0) as predictors. Given that *Rel Rew* ranges from -0.8 to 0.8, and that RT may be influenced by the similarity between options, we also included the absolute value of *Rel Rew* (|*Rel Rew*|).

### Computational Analysis

To quantify the dynamics of decision-making processes for approach-avoidance, we used the HDDM toolbox and applied different versions of Sequential Sampling Models that provided an algorithmic account of how evidence accumulation contributes to a binary approach vs. avoid decision (Fengler et al., 2021, 2022; Wiecki et al., 2013). Specifically, we applied simple drift diffusion model (DDM; Ratcliff, 1978), full-DDM (Ratcliff & McKoon, 2008), linear collapse model (LCM; Cisek et al., 2009), and Ornstein-Uhlenbeck model (OUM; Busemeyer & Townsend, 1993). Each model conceptualized that on each trial, participants may begin with an initial bias towards one decision and when the options were presented, they evaluated the presented options to accumulate evidence until there was sufficient information to reach a decision threshold. This process would be captured by four main parameters with well-established psychological significance and distinct effects on behavioral patterns: drift rate (v) that reflects how quickly and effectively information is integrated to reach a decision, boundary separation (a) that reflects the required amount of evidence for reaching decisions, starting point bias (z) that reflects the inherent bias towards a particular response boundary prior to observing the trial-specific contingencies, and non-decision time (t) that reflects time spent outside the decision process (e.g., perceptual encoding or motor execution). The full DDM includes additional inter-trial parameter variability (*sv, sz, st*); the OUM includes an additional leak (*g*) on drift rate such that the accumulated evidence gradually decays back towards a neutral point; and the LCM includes an additional collapse rate parameter (*θ*) that allows for the decision boundary to decline linearly over time.

The candidate models differed in how they represent the dynamics of evidence accumulation over time, and each model reflected distinct theoretical hypotheses about the decision-making process. All variants of these models allowed four main parameters to vary: (1) the rate at which evidence accumulated towards a decision (drift rate, *v*); (2) the amount of evidence required to make a decision (decision threshold, *a*; (3) a bias towards a particular response (starting point, *z*); and (4) elements of response time unrelated to the decision process (non-decision time, *t*). In addition, variants of the model allowed for the decision boundary to either remain static throughout the trial or to decline linearly over time (linear collapse model; LCM; Ging-Jehli, Kuhn et al., 2024).

For each model, we used hierarchical regression to test the impact of trial-by-trial variation in *Rel Pas*, *Rel Neg*, and *Cangruency* (binary coded as 0=incongruent trial and 1=congruent trial) on the decision parameters. Continuous model predictors were normalized before entering the regressions. Models were run with 3 chains, each with 10,000 samples (including 5,000 samples as burn-in). The Gelman-Rubin Ȓ statistic was used to assess model convergence (Ȓ<= 1.1 indicated model convergence; Gelman & Rubin, 1992). To assess the significance of the fitted parameters, we computed q-values by comparing the proportion of the Bayesian posterior distributions against 0.

Alternative modeling approaches that used Cumulative Prospect Theory models to investigate approach-avoidance decision dynamics in PAAT are reported in the Supplement. Nevertheless, the predictive performance and parameter recovery of the alternative models were worse compared to SSMs (Figure S4-6).

#### Model Validation and Selection

For each fitted model, we generated posterior predictive checks (PPC) to visualize potential misfits of the model by simultaneously considering response frequency and RT distribution for the responses. The best-fitting model was selected based on deviance information criterion (DIC) and PPC.

#### Parameter Recovery

After identifying and validating the best-fitting model, we performed simulation-based parameter recovery by sampling from normal distributions *N (pasteriar mean, pasteriar sd)* of the fitted parameters and fitting the model on synthetic data to determine whether the parameters can be recovered (Wiecki et al, 2013; Wilson & Collins, 2019).

## Results

To examine how sensitive approach-avoid decisions were to the likelihood of a given outcome, we parametrically varied the probabilities of rewarding and aversive outcomes between two options, one riskier (higher likelihood of a negative outcome) and one safer (lower likelihood of a negative outcome). To examine how these choice dynamics varied as a function of motivational congruency (i.e., the alignment between approach and avoid motivations), we also varied whether the safer option (lower likelihood of a negative outcome) was associated with a higher probability of a reward than the riskier option (motivationally *congruent*), or whether the safer option was associated with a lower probability of reward (motivationally *incongruent*). Collectively, this task enabled us to examine the influence of these varying outcome probabilities on the dynamics of these approach-avoid decisions.

### Model-Agnostic Analyses

#### Study 1

Focusing first on the choice participants made on incongruent trials (where rewarding and aversive outcomes supported divergent choices), mixed-effects regressions showed that participants were more likely to select the risky option the more it increased their chance of reward compared to the safer option (higher relative reward probability; B = 2.80, 95% CI = [1.84 3.76], p<.001). At the same time, participants were less likely to choose the riskier option the more it increased their chances of an aversive outcome (higher relative aversive probability; B = -0.97, 95% CI = [-1.24 -0.69], p<.001) (Figure 2). On congruent trials, neither relative reward probability nor relative aversive outcome probability influenced choices (ps > .10). Collapsing across all trials, we saw that participants were overall more likely to choose the safe option on congruent trials (when the safe option was more rewarding) compared to incongruent trials (when the safe option was less rewarding; B = -3.60, 95% CI = [-5.33 -1.86], p<.001; Table 2).

**Figure 2.**
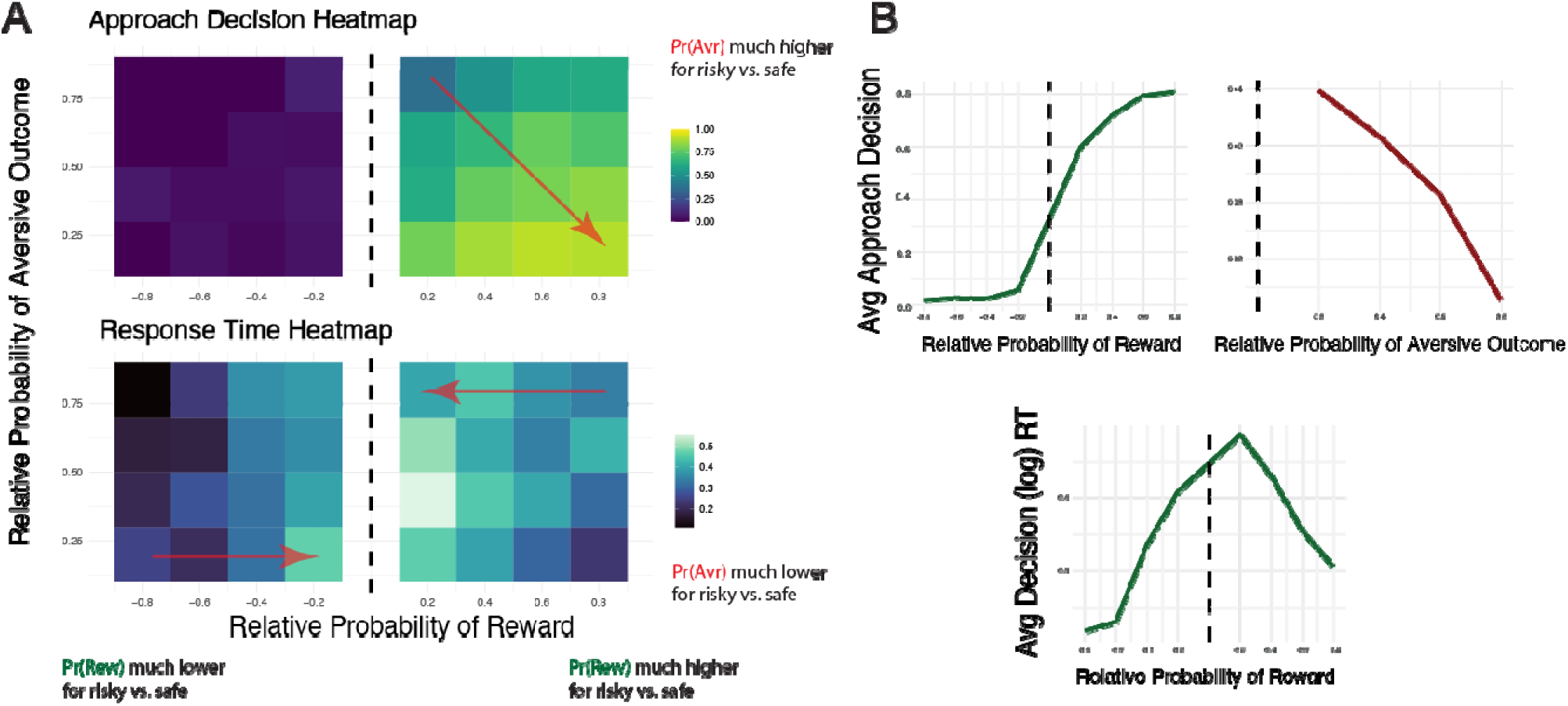
Study 1 behavioral patterns in PAAT. (A) Mean rates for choosing the risky option and average response times for presented congruent (with negative, left side of the dashed line) and incongruent trials (with positive, right side of the dashed line). In the choice heatmap (top panel), the red arrow indicated the trend of increasingly choosing the risky option as the increased and decreased. In the Response Time Heatmap (bottom panel), the red arrow indicated the trend of longer response time as the absolute value of decreased. (B) Mean rates for choosing the risky option as or varied, and average response times as varied. See behavioral patterns replications from Study 2 in Supplement Figure S7.

**Table 2.**
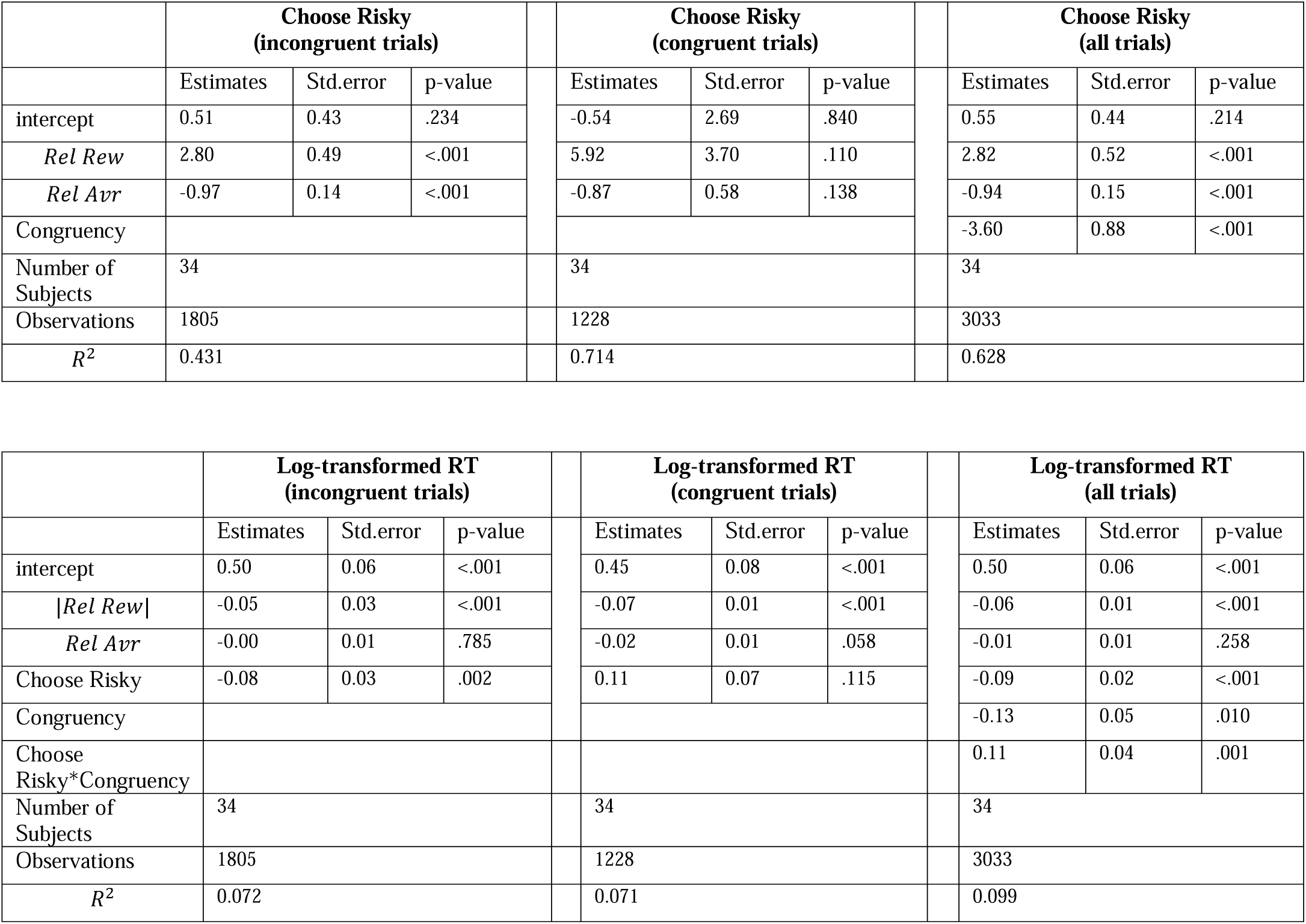
Study 1 mixed model results for approach choices and RT (log-transformed) using incongruent trials, congruent trials, and all trials. See Study 2 mixed model results in Supplement Table S7.

Response times revealed that participants were faster to choose on incongruent trials the greater the difference in reward likelihoods between the safer and riskier option (larger |*Rel Rew*|; B = -0.05, 95% CI = [-0.08 -0.03], p<.001). We did not find a comparable effect for negative outcome probabilities (*Rel Avr*; B = -0.00, 95% CI = [-0.02 0.02], p = .785) (Figure 2B). RTs also differed depending on which option they chose, such that participants responded faster when choosing the risky relative to the safe option (B = -0.08, 95% CI = [-0.13 -0.03], p =.002). On congruent trials, we observed similar patterns such that participants chose faster as the safer option increased the chance of reward compared to the risky option (B = -0.07, 95% CI = [-0.09 -0.05], p <.001). Collapsing across all trials, participants responded faster on congruent trials (B = -0.13, 95% CI = [-0.23 -0.03], p =0.010), and participants responded slower when choosing the risky option on congruent trials compared to incongruent trials (B *_*congruency*_* _X_ *_choice_* = 0.11, 95% CI = [0.05 0.18], p=.001; Table 2).

We also investigated how outcome probabilities for each option, individual differences in reactivity to IAPS images, and recent trial outcomes may influence task behavior (Supplement - Methods). Using the outcome probability of each option as predictors (instead of the relative values) found comparable effects, such that participants preferred an option (risky or safe option) as its trial-wise probabilities of rewarding outcome increased or trial-wise probabilities of aversive outcome decreased (Table S3). Neither recent trial history nor individual differences in IAPS image ratings influenced choice behavior or subjective estimates of outcome probabilities (Supplement Results - IAPS Images).

#### Study 2

Using data from an independent online sample, we replicated the influences of higher relative reward probability (B =1.50, 95% CI = [1.10 1.91], p<.001) and higher relative aversive probability (B = -0.44, 95% CI = [-0.61 -0.28], p<.001) on incongruent trials, as observed in Study 1 (which consists of a mixed clinical sample). On congruent trials, higher relative reward probability also influenced choices (B =1.51, 95% CI = [0.30 2.71], p=0.01). Similar to the findings from the discovery sample, participants were also more likely to choose the safe option on congruent trials compared to incongruent trials (B =-2.88, 95% CI = [-3.71 - 2.06], p<.001). We observed similar variations in RTs depending on difference in reward likelihoods and choices: participants were faster to choose the greater the difference in reward likelihoods between the safer and riskier option on incongruent (B = -0.33, 95% CI = [-0.43 - 0.24], p < .001) and congruent (B = -0.33, 95% CI = [-0.46 -0.19], p <.001) trials. Participants were faster when choosing the risky relative to the safe option on incongruent trials (B = -0.11, 95% CI = [-0.15 -0.08], p < .001), and were slower when choosing the risky option on congruent trials compared to incongruent trials (B *_congruency_* _X_ *_choice_* = 0.10, 95% CI = [0.05 0.15], p<.001; Table S7).

### Model-Based Analyses

We have thus far shown that variability in outcome probabilities, valence, and motivational congruency influence both the timing and outcomes of the decisions. Sequential sampling models (SSMs) enable the joint modeling of these two dependent variables (choices and RTs), to quantitatively capture changes in latent decision parameters across trials. To quantify the dynamics of approach-avoidance decision-making, we therefore fit SSMs to choice behavior in the PAAT task, with the aim of identifying the model parameterization that best accounts for these data.

#### Study 1

We found that the best-fitting model included trial-wise influences of relative reward probability, relative aversive probability, and their interaction with motivational congruency on drift rate (Table 3). Specifically, we found that, on incongruent trials, drift rate towards the risky option increased as a function of the relative probability of the rewarding outcome (B = 0.63, 95 % CI = [0.50 0.75], q-value <.001) and decreased as a function of the relative probability of the aversive outcome (B = -0.30, 95 % CI = [-0.37 -0.23], q-value <.001) (Figure 3). On congruent trials, drift rate increased with higher relative reward probability to a similar degree as on incongruent trials (B = 0.49, 95 % CI = [0.33 0.64], q-value <.001). By contrast, the negative effect of the relative aversive probability was smaller and not significant (B = -0.04, 95 % CI = [-0.11 0.03], q-value = .11). Across all trials, decisions were also characterized by a biased starting point for evidence accumulation, manifesting as a default bias towards choosing the risky option (*z*_*intercept*_ = 0.53, 95 % CI = [0.51 0.55], q-value <.001). Supplementary analysis using individual outcome probabilities (i.e., probability of positive and negative outcome associated with each option) found similar effects, such that drift rate towards an option (risky or safe) increased as the option’s probability of reward increased, or as the option’s probability of aversive outcome decreased (Figure S3).

**Figure 3.**
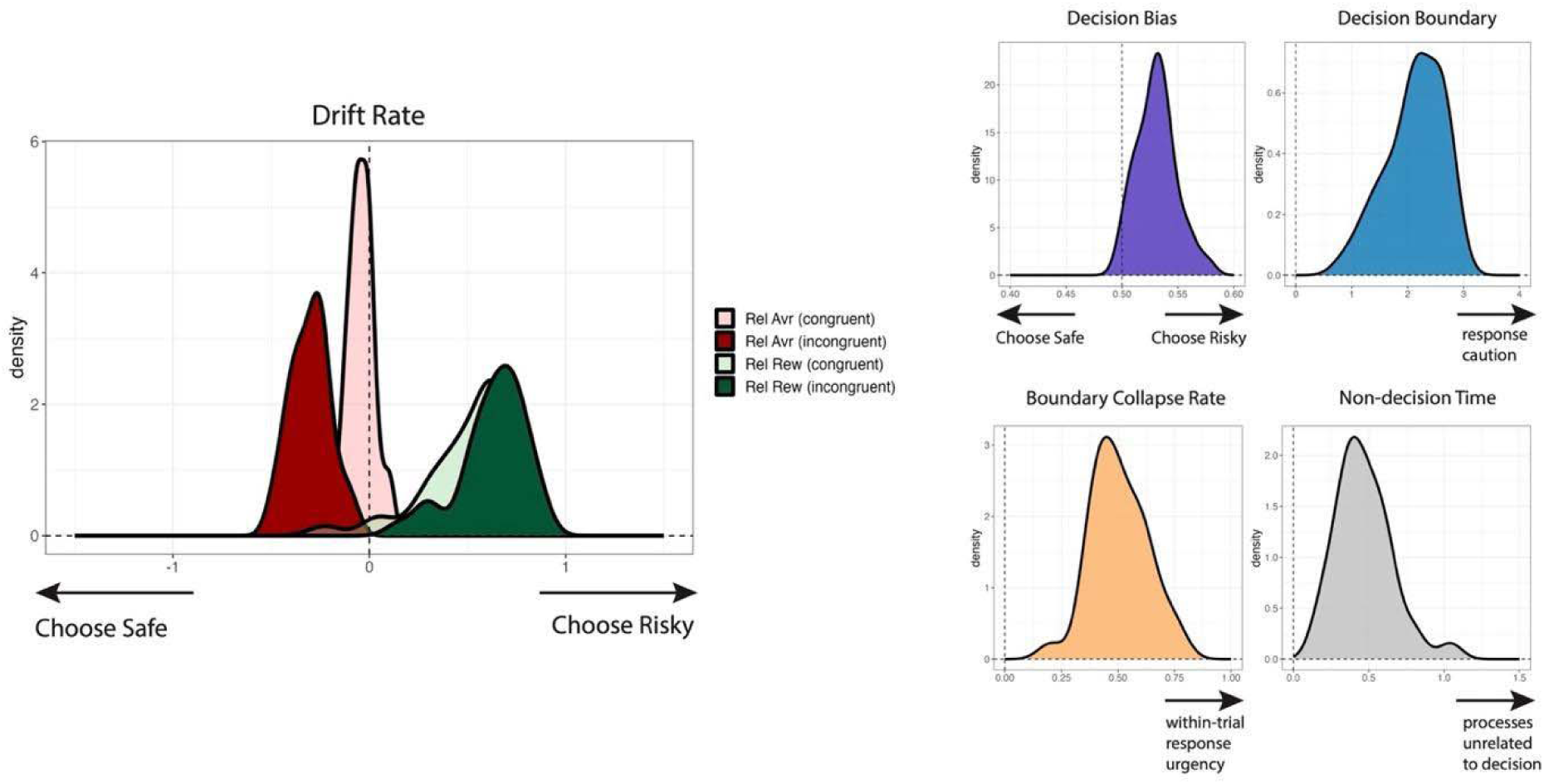
Study 1 group-level posterior distributions of the LCM parameters. Left: higher relative reward probability of the risky option influenced drift rate towards the risky option (on both incongruent and congruent trials; dark and light greens). Higher relative aversive outcome probability influenced drift rate towards the safe option (particularly on incongruent trials; dark red). Right: visualizations of the decision bias (purple), decision boundary (blue), boundary collapse rate (yellow), and non-decision time (grey) parameter distributions. See group-level posterior distributions of the LCM parameters replications from Study 2 in Supplement S8.

**Table 3.**
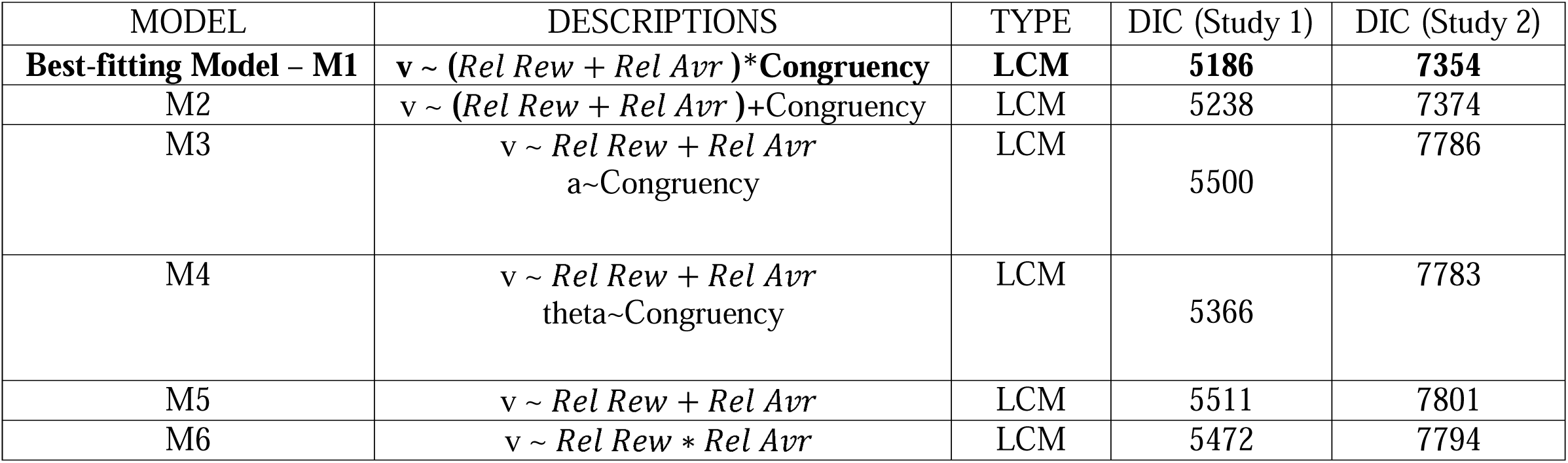
Different LCMs with the best-fitting model (selected based on DIC and PPC) represented in the first row. DIC refers to the deviance information criterion.

Our best-fitting model also incorporated a threshold that collapsed linearly over the course of a decision, but was not otherwise modulated by relative reward or aversive probability or motivational congruency (see Table S4 for model comparisons). This model provided a better fit to the data than equivalent models with static thresholds and/or additional parameters than can sometimes capture similar patterns in the data (e.g., models with leaks in evidence accumulation or with inter-trial parameter variability; Table S4). Test of model validation showed that the best-fitting model showed good posterior predictive performance and parameter recovery (Figure 4). For instance, simulated data captured the qualitative patterns seen on incongruent trials, whereby participants chose the risky option less frequently and had longer RTs for risky choices as conflict increased (filled red, blue, and green squares).

**Figure 4.**
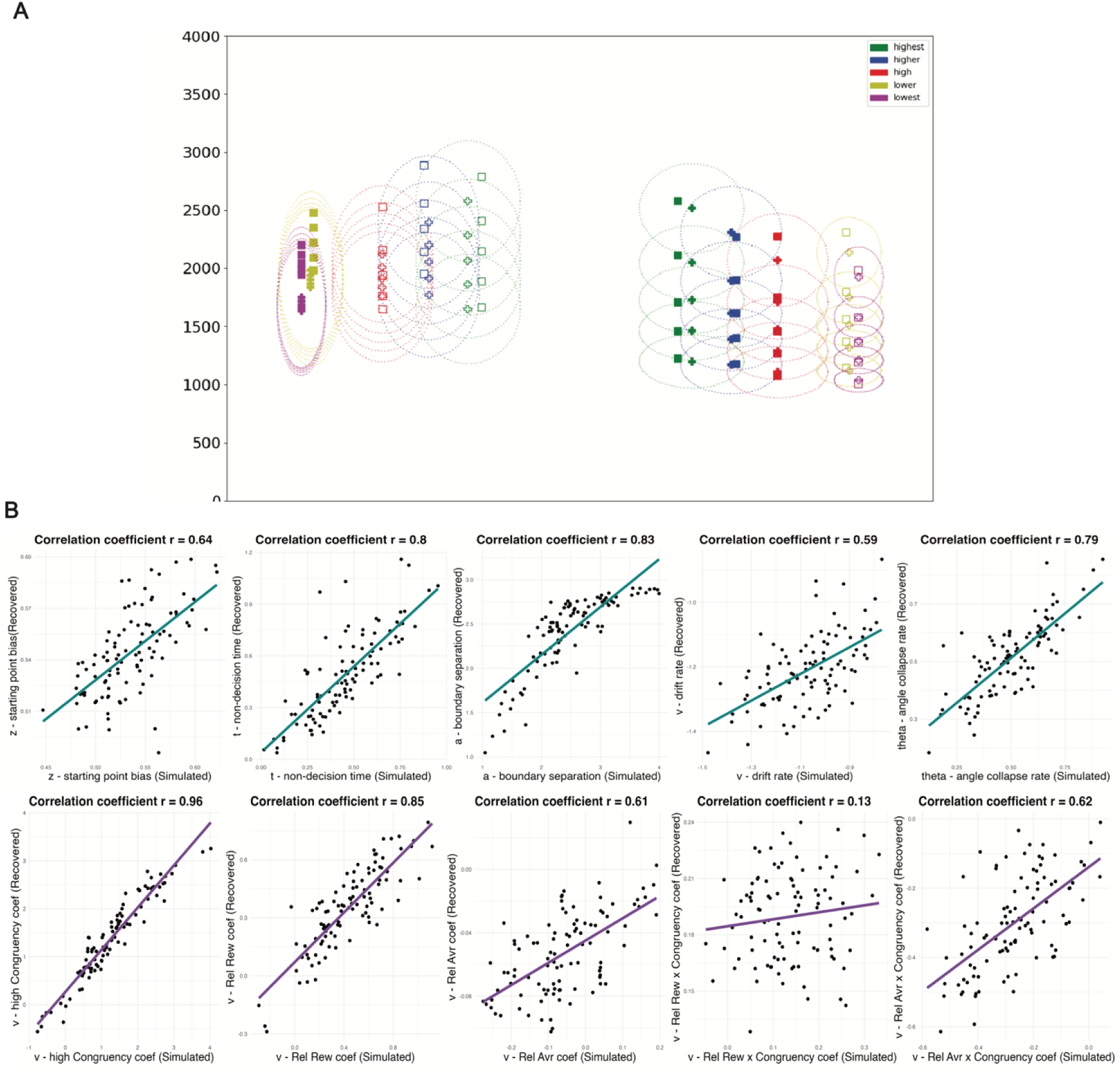
Study 1 Computational model validation. (A). Posterior predictive check of the best-fitting model. The x-axis represents choice frequency of choosing an option (filled shapes represent choosing the risky option and unfilled shapes represent choosing the safe option). Vertical columns moving from bottom to top represent the 0.1, 0.3, 0.5, 0.7, and 0.9 quantile response times. Empirical data are represented as squares and simulated data are represented as crosses. Ellipse widths (represented by dotted lines) index standard deviation of posterior predictive distributions and indicate model estimation uncertainty. Trials were equally binned (separately for congruent and incongruent trials) into five conditions (lowest, low conflict, high, higher, and highest conflict) based on the relative probability of positive outcome. (B). Simulation results comparing generative parameters (x-axis) with recovered parameters (y-axis). See computational model validation results replications from Study 2 in Supplement Figure S9.

#### Study 2

The best-fitting model from the discovery sample (Study 1, mixed clinical sample) also captured data from the replication sample (Study 2, online sample) well (Table 3). We found that relative probability of the rewarding outcome (B = 0.59, 95 % CI = [0.48 0.69], q-value <.001) and relative probability of the aversive outcome (B = -0.16, 95 % CI = [-0.20 - 0.11], q-value <.001) modulated drift rate towards the risky option on incongruent trials, and relative probability of the rewarding outcome also modulated drift rate towards the risky option on congruent trials (B = 0.40, 95 % CI = [0.27 0.52], q-value <.001). Similar to Study 1, model validations showed that the best-fitting model had good posterior predictive performance and parameter recovery (rs = 0.26-0.90; Figure S9).

## Discussion

The current study introduced a novel behavioral paradigm, the Probabilistic Approach-Avoidance Task (PAAT), that systematically manipulated outcome uncertainty and motivational conflict to explore how humans integrate probabilities of positive and negative outcomes during approach-avoid decision-making. We validated the paradigm by demonstrating that choice behavior in the PAAT is influenced by the probability of rewarding outcomes, the probability of aversive outcomes, and motivational congruency. We also leveraged a computational model of decision dynamics to quantify the impact of probabilistic outcomes on dissociable decision parameters. Results revealed that the default tendency to choose a risky or safe option varied across individuals. In addition, as outcome uncertainty varied, individuals adjusted evidence accumulation towards a risky or safe option to appropriately balance the drives to approach positive outcomes and avoid negative outcomes. Finally, we established that the computational model provided recoverable, subject-specific parameter estimates. Together, our model-based analysis of PAAT provided an algorithmic account of how evidence accumulation varies with outcome type and outcome uncertainty to contribute to approach-avoid decision making.

Our work builds on previous studies of valued-based risky decision-making and decision-making under approach-avoid conflict, which have shown that people are sensitive to relative value differences and gamble congruency, and the speed and direction of evidence accumulation vary with decision outcomes (Gelskov et al., 2015; Botvinik-Nezer et al., 2019; Peters & D’Esposito, 2020; Chen et al., 2022; Ironside et al., 2020; Pedersen et al., 2021; Ging-Jehli et al., 2024). We extend these findings to show that similar dynamics govern the influence of *uncertainty* about reward and aversive outcomes, particularly in cases where the two outcomes produce motivational conflict (incongruent trials). Across both samples (mixed clinical and online), participants on average displayed a default tendency towards approach before any trial-level option information was presented (i.e., starting point bias towards the riskier option). After the options were revealed, participants integrated information about potential outcomes by accumulating evidence: the relative probability of reward served as evidence supporting approach, while the relative probability of an aversive outcome supported avoidance. Although the present study was limited in examining associations between OCD symptoms and default approach tendencies or outcome integration, the modeling results provided an algorithmic account of how evidence accumulation processes contribute to approach-avoid decision-making under outcome uncertainty, with relevance for future clinical follow-up studies.

On congruent trials, where approach and avoidance motivation align, people tended to choose the most valuable option (the one with higher probability of rewarding outcome and lower probability of aversive outcome) on the vast majority of trials. In spite of these choices being near ceiling, response times revealed that decision-making still varied systematically with outcome likelihoods. Our model-based analyses demonstrated that the relative probability of positive outcomes still influenced drift rate parametrically and to a similar degree on these congruent trials as on incongruent trials. The influence of aversive outcomes on drift rate was directionally consistent with their influence on incongruent trials, but smaller and non-significant. It is possible that this sample was under-powered to detect the latter effect (particularly given that there were fewer congruent than incongruent trials), so it will be important to test whether this asymmetry emerges in follow-up work. If so, it may suggest that participants either adjust their weighting of negative outcomes on congruent trials, or that they sequentially consider positive outcomes prior to negative outcomes across all trials, a possibility that bears testing with measures of attention (e.g., eye tracking) and relevant variants of SSMs (e.g., Krajbich et al., 2010; Ting & Gluth, 2024). Previous studies including congruent and incongruent trials in approach-avoidance tasks demonstrated that these trial types were helpful in identifying effective and ineffective avoidance, and the associations between behavioral flexibility and psychiatric symptoms (Weaver et al., 2020). Our model comparisons likewise demonstrated improved fits for including parameters that quantify the influence of relative probabilities in both trial types relative to a model that assume an overall shift in evidence accumulation as trial type varied. Therefore, the parametric manipulation of outcome uncertainty in PAAT allowed for in-depth analysis of how individuals adjust their choices under outcome uncertainty in different motivational contexts.

The variability we observe in decision parameters during approach-avoid conflict offers valuable insights into potential individual differences in the decision dynamics under outcome uncertainty, and point toward potentially fruitful directions for understanding maladaptive behavioral patterns, such as persistent avoidance of potential unlikely negative outcomes at the expense of foregoing potential rewards. For example, obsessive-compulsive symptoms may be associated with biased responding (e.g., bias towards choosing safe) and/or biased evaluation on the PAAT (e.g., heightened sensitivity to potential aversive outcomes; Gillan et al., 2014; Luigjes et al., 2016); anxiety symptoms may be associated with heightened sensitivity to potential aversive outcomes (larger absolute regression coefficient of aversive outcome on drift rate; Perkins et al., 2020); and depressive symptoms may be associated with reduced sensitivity to potential rewards (smaller absolute regression coefficient of rewarding outcome on drift rate; Pedersen et al., 2021; Ging-Jehli et al., 2024). Thus, investigating the relationship between diverse symptom domains and computational parameters derived from the PAAT task represents a promising avenue for future research to understand flexible or inflexible approach-avoid decisions in response to uncertain rewarding vs. aversive consequences.

While the current provides critical validation for the PAAT task as a tool for modeling variability in decision dynamics based on choice behavior under approach-avoid conflict, sequential sampling models applied to our task may not generalize as a model across approach-avoid tasks (e.g., Bach, 2015; Zorowitz et al., 2019; Letkiewicz et al., 2023) and the source of evidence accumulation in the approach-avoid choices remains an open question. Therefore, critical next steps for future work will be to leverage neural activity measured during these decisions, as well as during the anticipation and outcome phases, to obtain a more comprehensive understanding of how individuals process the potential or actual occurrence of rewarding versus aversive outcomes (Critchley et al., 2001; Oldham et al., 2018; Ben-Zion & Levy, 2025). In addition, the current study was limited by the blinding diagnostic status in investigating the relationships between OCD diagnosis or symptom severity and approach-avoidance conflict behavior. Future studies in larger, clinically diverse samples may combine PAAT with neural circuit-level computational models to identify reliable neurocomputational biomarkers. Relatedly, while the current analyses focused on the relative differences in probability of rewarding or aversive outcome between the options, future studies can examine the impact of individual probability attributes (cf. Supplement Table S3), and the brain regions involved in signaling one or more of these variables. This approach would help link neural mechanisms with specific decision parameters and translate neuroscientific findings into a principled understanding of behavior and mental health (Montague et al., 2011; Wiecki et al., 2015; Huys et al., 2016).

## Authors’ contributions

Conceptualization: Ziwei Cheng, Nadja R. Ging-Jehli, Michael J. Frank, Mary L. Phillips, Amitai Shenhav

Data curation: Maisy Tarlow, Joonhwa Kim, Henry W. Chase, Manan Arora, Lisa Bonar, Ricki Stiffler, Alex Grattery, Simona Graur

Formal Analysis: Ziwei Cheng, Nadja R. Ging-Jehli

Supervision: Michael J. Frank, Mary L. Phillips, Amitai Shenhav

Writing – Original Draft Preparation: Ziwei Cheng, Nadja R. Ging-Jehli, Michael J. Frank, Amitai Shenhav

Writing – Review & Editing: all co-authors

## Declarations

## Acknowledgement

We are grateful to Ilya Monosov and Steve Rasmussen for helpful discussions and valuable input.

## Ethics approval

This research received approval from the Institutional Review Board of University of Pittsburgh.

## Consent to participate

All participants included in the study provided informed consent.

