## Supplemental_Materials for "A Novel Approach-Avoidance Task to Study Decision Making Under Outcome Uncertainty"

**Supplementary Methods**

***Exclusion criteria***

All participants

- Not meeting inclusion criteria
- Personal/family history (1st/2nd degree relative) of schizophrenia/schizoaffective disorder, other primary psychotic disorder, Bipolar Disorder
- Present PTSD, psychotic symptoms (SCID5 criteria), suicidal ideation (endorses yes to question 2 or higher on Columbia-Suicide Severity Rating Scale)
- Personal history of head injury, neurological disorder, neurodevelopmental disorder, tic disorder, and/or systemic medical diseases
- Intoxicated on scan day (salivary alcohol tests) and/or history in the last 3 months of illicit substance use and/or any substance use disorder (SUD), determined by scan day urine test for illicit substances, and SCID5 assessment of SUD
- MRI contraindications (metallic foreign objects, prone to panicking in enclosed spaces, positive pregnancy test/self-reporting pregnancy)
- Young Mania Rating Scale (YMRS) greater than or equal to 8

Healthy controls

- Adult onset/present Axis I disorders/ SUD
- Use of psychotropic medication
- Family history of schizophrenia/schizoaffective disorders, primary psychotic disorders and/or Bipolar Disorder
- Personal childhood history of psychiatric disorders (except childhood mood/anxiety disorders and ADHD are permitted)
- Hamilton Depression Rating Scale greater than or equal to 8

OCD participants

- Present Yale-Brown Obsessive Compulsive Scale (YBOCS) score less than 16
- OCD diagnosis with SCID 5 Hoarding Disorder
- Not meeting criteria for OCD diagnosis
- Taking psychotropic medications within the last 3 months (1 SRI or clomipramine is allowed)

***Inclusion criteria***

All participants

- 18-35 years of age
- Right-handedness (Annett criteria)
- Score 85 or higher on the North American National Adult Reading Test
- Score 24 or higher on the Mini-Mental State cognitive state examination
- Snellen visual acuity (>20/40)
- YMRS<8

Healthy controls

- No personal childhood history of psychiatric disorders (childhood mood/anxiety disorder history is permitted)
- No use of psychotropic medications
- No adult onset/present Axis I disorders
- Hamilton Depression Rating Scale <8

OCD participants

- Score 16 or higher on YBOCS
- OCD diagnosis without SCID 5 Hoarding Disorder
- Psychotropic medication free in the last 3 months or taking only one serotonin reuptake inhibitor (SRI) antidepressant medication or clomipramine

**Analysis with IAPS Images**

Before the start of the task, participants rated stimuli from the International Affective Picture System (25 negative and 25 neutral images). For each image, participants rated the extent to which the image made them feel unpleasant/negative (1) or pleasant/positive (7) on a 7-point scale (5 = neutral). They also rated the extent to which the image made them feel relaxed/calm (1) or stimulated/aroused (7) on the same scale.

We conducted exploratory analysis using Study 1 data to account for the possibility that participants’ task behavior will be influenced by individual differences in reactivity to the IAPS images. For each participant, we characterized individual reactivity to the images as the difference between their average valence and arousal ratings between the negative and neutral images

$R_{val}=Valence \left( negative \right)$ - $Valence \left( neutral \right)$

$R_{arl}=Arousal \left( negative \right)$ - $Arousal (neutral)$)

To investigate whether individuals who (on average) rated the negative images as more unpleasant or arousing compared to neutral images showed different choice behavior, we included $R_{val}$ and $R_{arl}$ as additional predictors in a regression models to predict approach decisions (choose the risky option).

$Choose risky \sim\left( Rel Rew+Rel Avr \right)*(R_{val}$ + $R_{arl}$) + Congruency

**Analysis with Recent Trial Outcome**

To account for the potential influence of recent trial outcomes (reward, neutral, or negative image) on behavior, we used Study 1 data and tested whether the previous trial’s outcome would predict decisions to choose risky on the current trial in the regression models.

$Choose risky \sim\left( Rel Rew+Rel Avr \right)*PrevOutcome$ + Congruency

We also tested whether the trial history would alter the default likelihood of choosing the risky vs. safe option in the SSMs (reflected in modulation of starting point, z).

**Analysis with Individual Reward and Aversiveness of the Options**

The main analysis focused on relative probability of rewarding outcome and aversive outcome between the risky and safe options, and we note that we can also model the influence of individual probability of rewarding and aversive outcomes (P${r\left( Rew \right)}_{risky}$, $Pr\left( Rew \right)_{safe}$,$P{r\left( Avr \right)}_{risky}$, and $Pr\left( Avr \right)_{safe}$) on the decision dynamics. Although the relative values captured variability in behavior across PAAT trials with fewer parameters (Table S4, S5), using the trial-wise, individual probability of rewarding and aversive outcomes may be useful for future studies identifying brain regions involved in signaling these task variables. Therefore, we used Study 1 data and tested regression models and SSMs with individual probability of rewarding and aversive outcomes to validate the paradigm manipulations.

**Analysis with** **Cumulative Prospect Theory Models**

We explored the Cumulative Prospect Theory (CPT) model, a prominent model of decision-making (Tversky & Kahneman, 1992), as alternative models for understanding the mental processes underlying evaluations of different outcome probabilities in PAAT. To evaluate the feasibility and utility of fitting CPT to the PAAT, we used Study 1 data and implemented multiple CPT model variants to (1) evaluate model fit, (2) evaluate model recovery, and (3) evaluate parameter recoverability.

***Model 1: Full CPT***

On each trial, the model evaluates the objective probabilities in an option with a probability weighting function:

$w\left( p; \gamma\right)=\frac{p^{\gamma}}{\left[ p^{\gamma}+ \left( 1 - p \right)^{\gamma} \right]^{\frac{1}{\gamma}}}$	Equation S1

$\gamma$ is separately estimated for evaluating the probability of a positive outcome ($\gamma_{pos}$), and for evaluating the probability of a negative outcome ($\gamma_{neg}$). The magnitudes of gain and loss outcomes were fixed to 5 units. To capture individual differences in weighing the different outcomes, the objective outcome magnitude was transformed by a power utility function with exponents α (for gains) and β (for losses). For each option, the subjective value (utility) was computed as Equation S2, where λ represents loss aversion:

$V = w\left( p_{pos}; \gamma_{pos} \right)* 5^{\alpha}- \lambda* w\left( p_{neg}; \gamma_{neg} \right)* 5^{\beta},$	Equation S2

The model compares the utility difference between the options, and the probability of choosing one option over the other (e.g., choose right vs. left) was calculated as Equation S3, with θ representing a sensitivity parameter (inverse temperature) that governs the determinism of choice behavior:

$P\left( choose right \right)= \frac{1}{\left[ 1 + exp\left( -\theta* \left( V_{right}- V_{left} \right) \right) \right]}$	Equation S3

Together, this model included six parameters that can be fit to each participant’s data: $\gamma_{pos}$ (sensitivity to positive outcome probability), $\gamma_{neg}$ (sensitivity to negative outcome probability), $\alpha$ (weighing of positive outcome), $\beta$ (weighing of negative outcome), λ (loss aversion), and θ (choice stochasticity).

***Model 2: Simplified CPT with 4 parameters (SCPT-4)***

Since the magnitude of outcomes did not change in the task and parameters that estimate weighing of different outcomes may trade off with parameters that estimate weighing of different outcome probabilities, we fitted a simplified CPT with 4 parameters: $\gamma_{pos}$ (sensitivity to positive outcome probability), $\gamma_{neg}$ (sensitivity to negative outcome probability), λ (loss aversion), and θ (choice stochasticity).

***Model 3: Simplified CPT with 4 parameters (SCPT-3)***

Since loss aversion may be captured by comparing $\gamma_{neg}$ to $\gamma_{pos}$ (individuals who are aversive to losses would be more sensitive to potential negative outcomes), we fitted a simplified CPT with 3 parameters: $\gamma_{pos}$ (sensitivity to positive outcome probability), $\gamma_{neg}$ (sensitivity to negative outcome probability), and θ (choice stochasticity).

**Model Fitting**

Parameters were estimated using maximum likelihood estimation. We used the L-BFGS-B algorithm, implemented by *scipy.optimize.minimize* in Python (Virtanen et al., 2020). $\gamma_{pos}$, $\gamma_{neg}$, $\alpha$, and $\beta$ ranged between 0 and 1, and λ (loss aversion), and θ (choice stochasticity) ranged between 0 and 2.

**Supplementary Results**

**IAPS Images**

Participants found negative images to be less pleasant (B = -2.35, p<.001) and more arousing (B = 1.97, p <.001) than neutral images. Individuals who rated the negative images as more unpleasant or more arousing than the neutral images did not differ from others in their frequencies to choose the risky option (nonsignificant main effects of $R_{val}$ and $R_{ars}$; ps > .56), nor in their subjective estimates of the outcome probabilities when making decisions during the task (nonsignificant interactions between $R_{val}$ and $R_{ars}$, and $Rel Rew$ and $Rel Avr$, ps > .27).

In the regression models, we did not find any main effect of previous trial’s outcome on decisions (B_pos_ = 0.01, p = 0.98, B_neg_ = -0.86, p = 0.07), nor any interactions between previous trial’s outcome and trial-level task manipulation (nonsignificant interactions between $PrevOutcome$, and $Rel Rew$ and $Rel Avr$, ps > .23). In the sequential sampling model, we did not observe any effect of the previous trial’s outcome on decision bias (B_pos_ = 0.01, 95 % CI = [-0.01, 0.03]; B_neg_ = 0.01, 95 % CI = [-0.01, 0.03]; ps > .05).

**Individual Reward and Aversiveness of the Options**

Models in the main text simplified the task variables into two relative values ($Rel Rew$ and $Rel Avr$), whereas complementary analyses examined the influence of individual probabilities of rewarding and aversive outcomes on decision dynamics (Table S3). Regression analysis including congruency and individual outcome probabilities of risky and safe options found that on incongruent trials, participants were more likely to choose risky as the probability of rewarding outcome of the risky option increased (B = 1.87, p < .001) or as the probability of aversive outcome of the safe option increased (B = 0.82, p < .001). Participants were less likely to choose risky as the probability of aversive outcome of the risky option increased (B = -1.22, p < .001) or as the probability of rewarding outcome of the safe option increased (B = -2.16, p < .001). Compared to incongruent trials, participants were more likely to choose the safe option over the risky option on congruent trials (B = -4.03, p < .001).

We also applied a sequential sampling model with linear collapsing boundaries to test the influence of individual outcome probability of the options on the decision parameters. The results showed that drift rate towards choosing the risky option were higher in incongruent trials (B = 1.31, 95 % CI = [0.91, 1.71], p <.001), were higher as the probability of negative outcome of the safe option increases (B = 0.19, 95 % CI = [0.13, 0.25], p < .001), and were higher as the probability of rewarding outcome of the risky option increases (B = 0.37, 95 % CI = [0.30, 0.43], p <.001). In addition, drift rate towards choosing the risky option were lower as the probability of rewarding outcome of the safe option increases (B = -0.38, 95 % CI = [-0.46, -0.29], p<.001) or the probability of negative outcome of the risky option increases (B = -0.23, 95 % CI = [-0.29, -0.16], p<.001).

Together, the results validated the paradigm manipulations by showing that higher probability of positive outcomes (of risky or safe option) pulled participants towards that option, and higher probability of negative outcomes pushed participants away from that option.

**Cumulative Prospect Theory Models**

Log-likelihood comparisons found that the three fitted models explained the data better than a null model that makes random choices (Figure S4). Among the three fitted models, the simplest model (SCPT3) had better fits (AIC or BIC) than the other ones. Although the CPT models can capture qualitative patterns in the data (Figure S5), model recovery and parameter recovery were poor among the CPT models applied to PAAT (Table S6; Figure S6).

|  | N = 34 | N = 50 |
| --- | --- | --- |
| **Gender** |  |  |
| Female | 24 (70.6%) | 23 (46%) |
| Male | 9 (26.4%) | 24 (48%) |
| Transgender | 1 (3%) | 0 (0%) |
| Other or prefer not to answer | 0 (0%) | 3 (6%) |
| **Age** | 25.2 (4.5) | 37.1 (10.1) |
| **Ethnicity** |  |  |
| Hispanic or Latino | 3 (9%) | 6 (12%) |
| Non-Hispanic and non-Latino | 31 (91%) | 44 (88%) |
| **Race** |  |  |
| African American | 2 (6%) | 5 (10%) |
| American Indian/Alaskan native | 0 (0%) | 0 (0%) |
| Asian | 7 (20.5%) | 4 (8%) |
| More than one race | 0 (0%) | 4 (8%) |
| Native Hawaiian/other Pacific Islander | 0 (0%) | 1 (2%) |
| White | 25 (73.5%) | 34 (68%) |
| Other or prefer not to answer | 0 (0%) | 2 (4%) |

Table S1. Demographics Information of the Sample in Study 1 and Study 2.

|  | **Choose Risky**  **(incongruent trials)** | | **Choose Risky**  **(congruent trials)** | | **Choose Risky**  **(incongruent trials)** | | **Choose Risky**  **(congruent trials)** | |
| --- | --- | --- | --- | --- | --- | --- | --- | --- |
|  | Estimates | p-value | Estimates | p-value | Estimates | p-value | Estimates | p-value |
| intercept | 0.76 | .77 | -0.55 | .95 | 0.82 | .78 | -4.08 | .52 |
| $Rel Rew$ | 6.34 | .01 | 12.75 | .222 |  | |  | |
| $Rel Avr$ | -2.17 | .01 | -5.17 | .06 |  |  |  |  |
| $Rel Rew* r_{incongruent}$ | -5.98 | .10 | -12.38 | .43 |  |  |  |  |
| $Rel Avr * r_{incongruent}$ | 1.97 | .13 | 7.31 | .07 |  |  |  |  |
| $r_{incongruent}$ | -0.35 | .93 | -0.62 | .96 | -0.25 | .96 | 0.18 | .99 |
| $Rew_{risky}$ |  | |  | | 4.60 | .01 | 7.18 | .13 |
| $Avr_{risky}$ |  |  |  |  | -3.34 | .01 | -5.63 | .06 |
| $Rew_{safer}$ |  |  |  |  | -5.08 | .02 | -5.14 | .31 |
| $Avr_{safer}$ |  |  |  |  | 2.00 | .04 | 3.59 | .26 |
| $Rew_{risky} * r_{incongruent}$ |  |  |  |  | -4.55 | .07 | -8.90 | .21 |
| $Avr_{risky} * r_{incongruent}$ |  |  |  |  | 3.41 | .07 | 8.51 | .06 |
| $Rew_{safer}* r_{incongruent}$ |  |  |  |  | 4.80 | .16 | 6.79 | .37 |
| $Avr_{safer}* r_{incongruent}$ |  |  |  |  | -1.76 | .24 | -6.63 | .17 |
| Number of Subjects | 34 | | 34 | | 34 | | 34 | |
| Observations | 1805 | | 1228 | | 1228 | | 1805 | |
| $R^{2}$ | 0.45 | | 0.76 | | 0.17 | | 0.59 | |

Table S2. Study 1 mixed model results including interactions between task variables and the ratio of incongruent trials for each participant.

|  | **Choose Risky**  **(incongruent trials)** | | |  | **Choose Risky**  **(congruent trials)** | | |  | **Choose Risky**  **(all trials)** | | |
| --- | --- | --- | --- | --- | --- | --- | --- | --- | --- | --- | --- |
|  | Estimates | Std.error | p-value |  | Estimates | Std.error | p-value |  | Estimates | Std.error | p-value |
| intercept | 0.66 | 0.47 | .158 |  | -2.03 | 2.77 | .464 |  | 0.66 | 0.48 | .172 |
| $Rew_{risky}$ | 1.82 | 0.34 | <.001 |  | 4.55 | 2.43 | .061 |  | 1.87 | 0.36 | <.001 |
| $Avr_{risky}$ | -1.25 | 0.20 | <.001 |  | -1.28 | 0.77 | .097 |  | -1.22 | 0.20 | <.001 |
| $Rew_{safer}$ | -2.15 | 0.41 | <.001 |  | -3.28 | 2.48 | .186 |  | -2.16 | 0.43 | <.001 |
| $Avr_{safer}$ | 0.91 | 0.17 | <.001 |  | 0.17 | 0.82 | .836 |  | 0.82 | 0.17 | <.001 |
| Congruency |  | | |  |  | | |  | -4.03 | 0.96 | <.001 |
| Number of Subjects | 34 | | |  | 34 | | |  | 34 | | |
| Observations | 1805 | | |  | 1228 | | |  | 3033 | | |
| $R^{2}$ | 0.457 | | |  | 0.725 | | |  | 0.881 | | |

Table S3. Study 1 mixed model results with individual reward and aversiveness of the options as predictors for approach choices using incongruent trials, congruent trials, and all trials.

| MODEL | DESCRIPTIONS | BIC (Study 1) | BIC (Study 2) |
| --- | --- | --- | --- |
| RM_Choice_1 | *ChooseRisky* ~ $Rel Rew$ + $Rel Avr$ *+ Congruency* | 1561 | 2881 |
| RM_Choice_S2 | *ChooseRisky* ~ $Rel Rew$ * $Rel Avr$ | 1569 | 2942 |
| RM_Choice_S3 | *ChooseRisky* ~ ($Rel Rew$ + $Rel Avr$*) * Congruency* | 1613 | 2983 |
| RM_Choice_S4 | *ChooseRisky* ~ ${Rew}_{risky}+{Rew}_{safe}+{Avr}_{risky}+{Avr}_{safe}$ + *Congruency* | 1638 | 2957 |
| RM_Choice_S5 | *ChooseRisky* ~ $Rel Rew$ * $Rel Avr$ *+ Congruency* | 1571 | 2910 |
| RM_Choice_S6 | *ChooseRisky* ~ $Rel Rew$ + $Rel Avr$ *+ Congruency +Avg Rew + Avg Avr* | 1607 | 2957 |
| RM_RT _1 | logRT ~$｜Rel Rew \vert$ + $Rel Avr$ *+ Congruency * ChooseRisky* | 2230 | 4220 |
| RM_RT _S2 | logRT ~ ($Rel Rew$ + $Rel Avr$ *+ ChooseRisky*) ** Congruency* | 2381 | 4374 |
| RM_RT _S3 | logRT ~ $Rel Rew$ + $Rel Avr$ *+ Congruency + ChooseRisky* | 2341 | 4410 |
| RM_RT _S4 | logRT ~ $Rel Rew$ * $Rel Avr$ *+ Congruency + ChooseRisky* | 2403 | 4471 |

Table S4. Comparisons between linear mixed-effects regression models used to analyze trial-level approach decisions (Choose the risky option) and response times (logRT) in the study. $Avg Rew= \frac{( {Pr\left( Rew \right)}_{risky}+ Pr\left( Rew \right)_{safe})}{2}, Avg Avr=\frac{P{r\left( Avr \right)}_{risky}+Pr\left( Avr \right)_{safer}}{2}$*.* Lower BIC values indicate better model fit.

| MODEL | DESCRIPTIONS | TYPE | DIC |
| --- | --- | --- | --- |
| **SSM1** | **v ~ (**$Rel Rew$ + $Rel Avr$ **)*Congruency** | **LCM** | **5186** |
| SSM_S2 | v ~ ${Rew}_{risky}+{Rew}_{safe}+{Avr}_{risky}+{Avr}_{safe}$+Congruency | LCM | 5233 |
| SSM_S3 | v ~ $Rel Rew$ + $Rel Avr$  *theta ~ EV difference * Congruency* | LCM | 5473 |
| SSM_S4 | v ~ ($Rel Rew$ + $Rel Avr$ *) * Congruency*  *theta ~ EV difference* | LCM | 5201 |
| SSM_S5 | v ~ ($Rel Rew$ + $Rel Avr$ *) * EV difference*  *theta ~ Congruency* | LCM | 5366 |
| SSM_S6 | v ~ $Rel Rew$ ***** $Rel Avr$ | OUM | 6117 |
| SSM_S7 | v ~ $Rel Rew$ ***** $Rel Avr$ | DDM | 5688 |
| SSM_S8 | v ~ $Rel Rew$ ***** $Rel Avr$ | Full-DDM | 5501 |

Table S5. Study 1 additional SSMs with the best-fitting model reported in main text as the first row. LCM: linear collapse model; OUM: Ornstein-Uhlenbeck model; DDM: drift diffusion model; Full-DDM: drift diffusion model with inter-trial parameter variability. EV difference was calculated as $| [{Pr\left( Rew \right)}_{risky}, - Pr\left( Rew \right)_{safe}]-[P{r\left( Avr \right)}_{risky}-Pr\left( Avr \right)_{safer}] |$ and represented a continuous measure of conflict (higher EV difference represented lower conflict, and lower EV difference represent higher conflict.

|  | CPT | SCPT 4 | SCPT 3 |
| --- | --- | --- | --- |
| $\gamma_{pos}$ | 0.38 | 0.89 | 0.87 |
| $\gamma_{neg}$ | 0.6 | 0.66 | 0.93 |
| $\alpha$ | 0.38 | - | - |
| $\beta$ | 0.39 | - | - |
| λ | 0.37 | 0.29 (n.s.) | - |
| θ | 0.87 | 0.01 (n.s.) | 0.05 (n.s.) |

Table S6: Study 1 CPT model parameter recovery summary. Recoverability was measured with the Pearson-correlation coefficient between true parameter and recovered parameter.

|  | **Choose Risky**  **(incongruent trials)** | | |  | **Choose Risky**  **(congruent trials)** | | |  | **Choose Risky**  **(all trials)** | | |
| --- | --- | --- | --- | --- | --- | --- | --- | --- | --- | --- | --- |
|  | Estimates | Std.error | p-value |  | Estimates | Std.error | p-value |  | Estimates | Std.error | p-value |
| intercept | 1.20 | 0.31 | <.001 |  | -1.76 | 0.68 | .10 |  | 1.18 | 0.31 | <.001 |
| $Rel Rew$ | 1.50 | 0.20 | <.001 |  | 1.51 | 0.62 | .014 |  | 1.64 | 0.20 | <.001 |
| $Rel Avr$ | -0.44 | 0.08 | <.001 |  | -0.30 | 0.25 | .226 |  | -0.46 | 0.08 | <.001 |
| Congruency |  | | |  |  | | |  | -2.88 | 0.42 | <.001 |
| Number of Subjects | 50 | | |  | 50 | | |  | 50 | | |
| Observations | 3264 | | |  | 1491 | | |  | 4755 | | |
| $R^{2}$ | 0.17 | | |  | 0.16 | | |  | 0.72 | | |

|  | **Choose Risky**  **(incongruent trials)** | | |  | **Choose Risky**  **(congruent trials)** | | |  | **Choose Risky**  **(all trials)** | | |
| --- | --- | --- | --- | --- | --- | --- | --- | --- | --- | --- | --- |
|  | Estimates | Std.error | p-value |  | Estimates | Std.error | p-value |  | Estimates | Std.error | p-value |
| intercept | 1.32 | 0.34 | <.001 |  | -1.77 | 0.78 | .024 |  | 1.23 | 0.33 | <.001 |
| $Rew_{risky}$ | 1.00 | 0.15 | <.001 |  | 1.62 | 0.55 | .003 |  | 1.18 | 0.15 | <.001 |
| $Avr_{risky}$ | -0.65 | 0.12 | <.001 |  | -0.22 | 0.33 | .512 |  | -0.65 | 0.11 | <.001 |
| $Rew_{safer}$ | -1.09 | 0.17 | <.001 |  | -0.85 | 0.50 | .087 |  | -1.07 | 0.17 | <.001 |
| $Avr_{safer}$ | 0.32 | 0.10 | .001 |  | 0.33 | 0.28 | .241 |  | 0.32 | 0.09 | <.001 |
| Congruency |  | | |  |  | | |  | -3.06 | 0.46 | <.001 |
| Number of Subjects | 50 | | |  | 50 | | |  | 50 | | |
| Observations | 3264 | | |  | 1491 | | |  | 4755 | | |
| $R^{2}$ | 0.19 | | |  | 0.23 | | |  | 0.73 | | |

|  | **Log-transformed RT**  **(incongruent trials)** | | |  | **Log-transformed RT**  **(congruent trials)** | | |  | **Log-transformed RT**  **(all trials)** | | |
| --- | --- | --- | --- | --- | --- | --- | --- | --- | --- | --- | --- |
|  | Estimates | Std.error | p-value |  | Estimates | Std.error | p-value |  | Estimates | Std.error | p-value |
| intercept | 0.39 | 0.05 | <.001 |  | 0.32 | 0.06 | <.001 |  | 0.38 | 0.05 | <.001 |
| $\vert Rel Rew\vert$ | -0.33 | 0.05 | <.001 |  | -0.33 | 0.07 | <.001 |  | -0.33 | 0.05 | <.001 |
| $Rel Avr$ | -0.00 | 0.01 | .524 |  | -0.01 | 0.01 | .190 |  | -0.01 | 0.01 | .227 |
| Choose Risky | -0.11 | 0.02 | <.001 |  | 0.04 | 0.04 | .312 |  | -0.10 | 0.01 | <.001 |
| Congruency |  | | |  |  | | |  | -0.11 | 0.03 | 0.001 |
| Choose Risky*Congruency |  | | |  |  | | |  | 0.10 | 0.03 | <.001 |
| Number of Subjects | 50 | | |  | 50 | | |  | 50 | | |
| Observations | 3264 | | |  | 1491 | | |  | 4755 | | |
| $R^{2}$ | 0.10 | | |  | 0.02 | | |  | 0.08 | | |

Table S7. Study 2 mixed model results for approach choices and RT (log-transformed) using incongruent trials, congruent trials, and all trials.

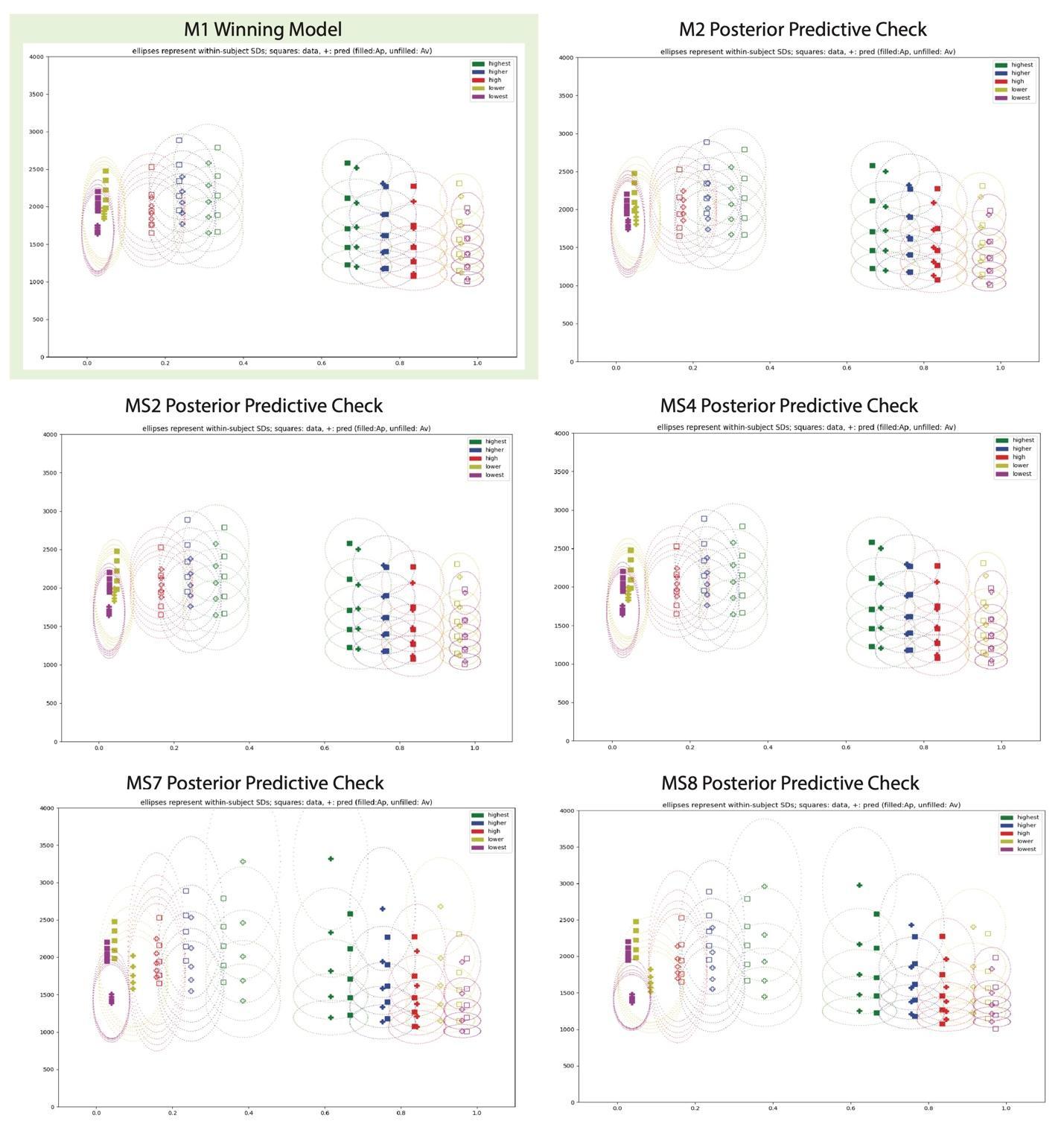

Figure S1. Study 1 posterior predictive check of selected alternative models (compared to best-fitting model) reported in manuscript and Table S5. In congruent trials (lowest and low conflict), participants almost always chose the safe option. In incongruent trials, participants chose the risky option less frequently and had longer RTs when they chose the risky as conflict increased (filled red, blue, and green squares). The winning model reported in the manuscript successfully captured the qualitative patterns.

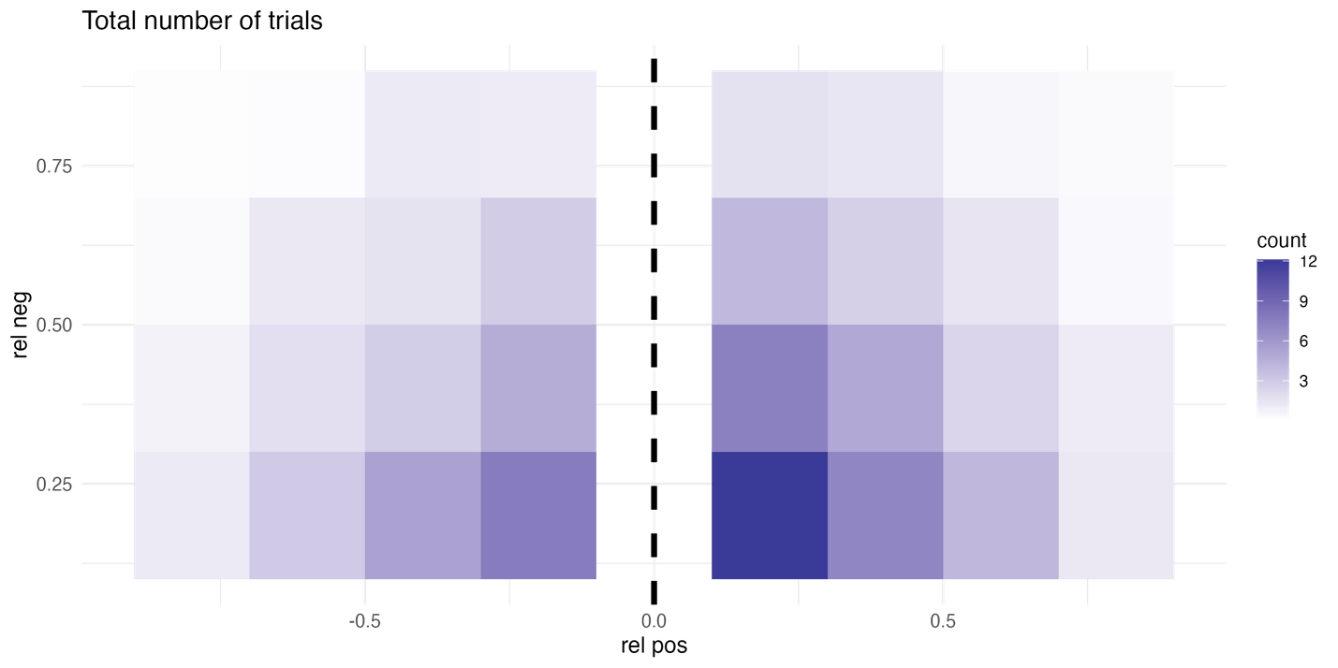

Figure S2. Study 1 number of trials for presented congruent and incongruent trials (averaged across all participants).

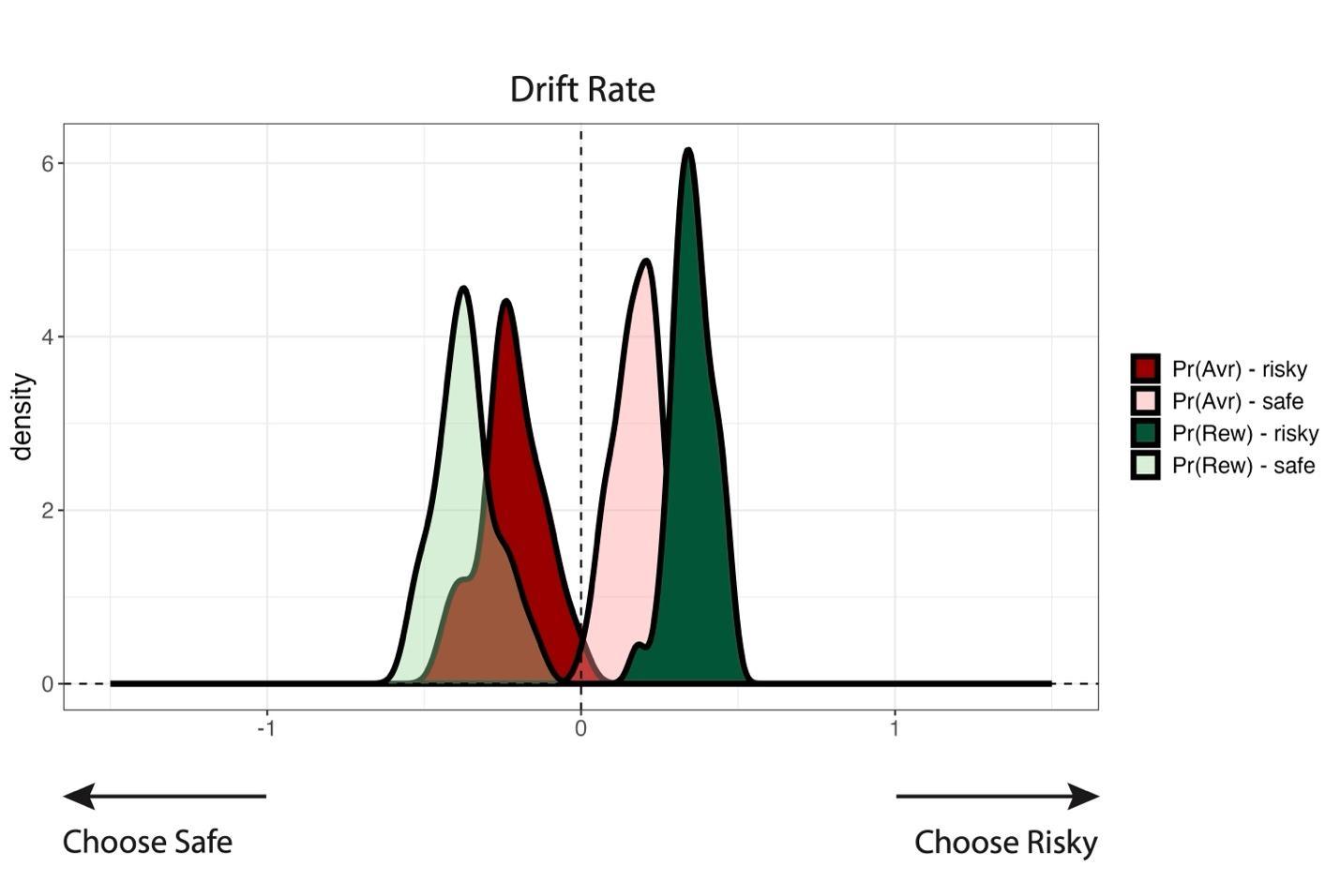

Figure S3. Study 1 posterior distributions of the LCM parameters from a model with individual outcome probability of the risky and safe options.

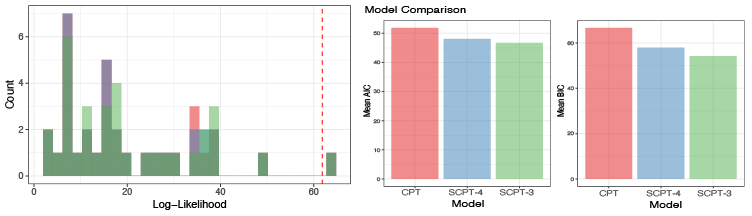

Figure S4. Study 1 CPT model comparison results. Left: comparisons between the three fitted models and a null model that makes random choices (likelihood at dashed line). Right: model comparisons (AIC or BIC) between the three fitted models

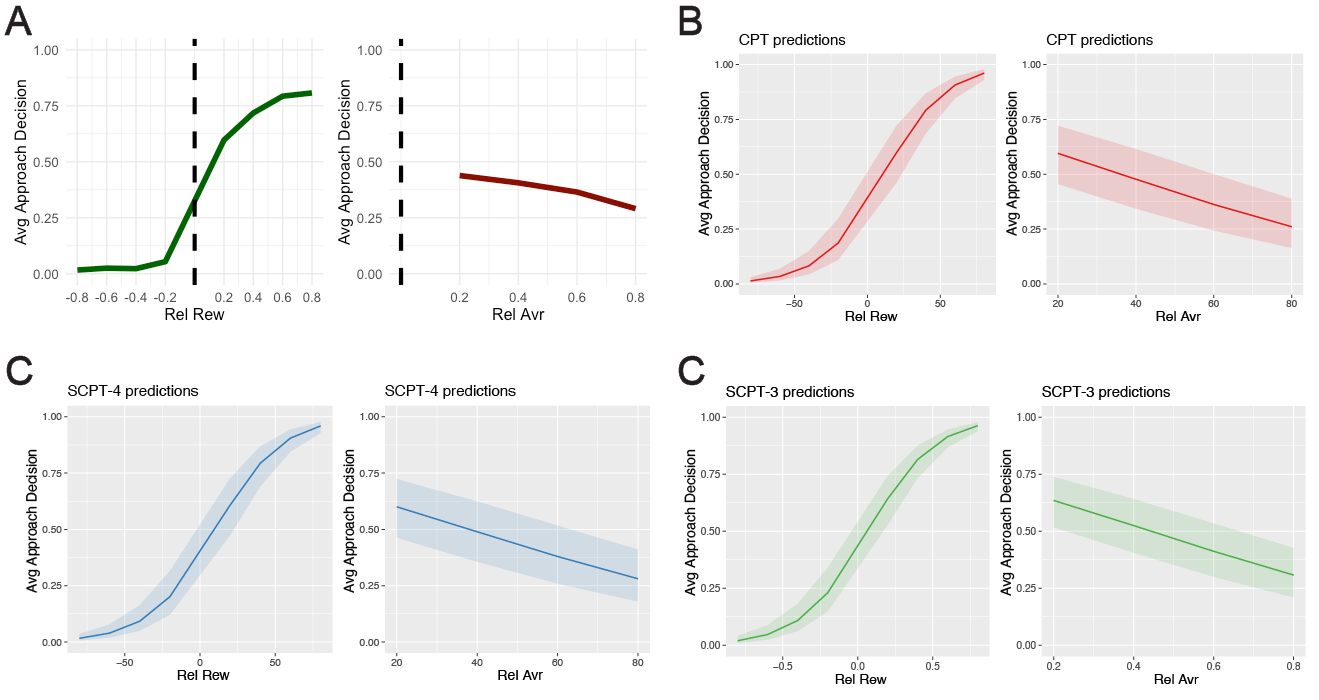

Figure S5. Study 1 CPT Model predictive performance. (A). summarized qualitative patterns of how decisions varied with $Rel Rew$ and $Rel Avr$ in the data. (B-D): summarized simulated patterns from each model.

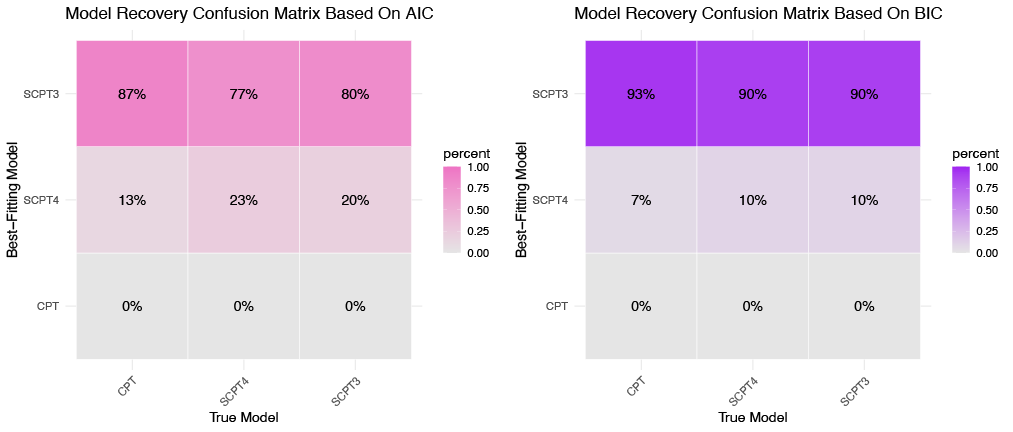

Figure S6. Study 1 CPT model recovery using AIC (Left) and BIC (Right).

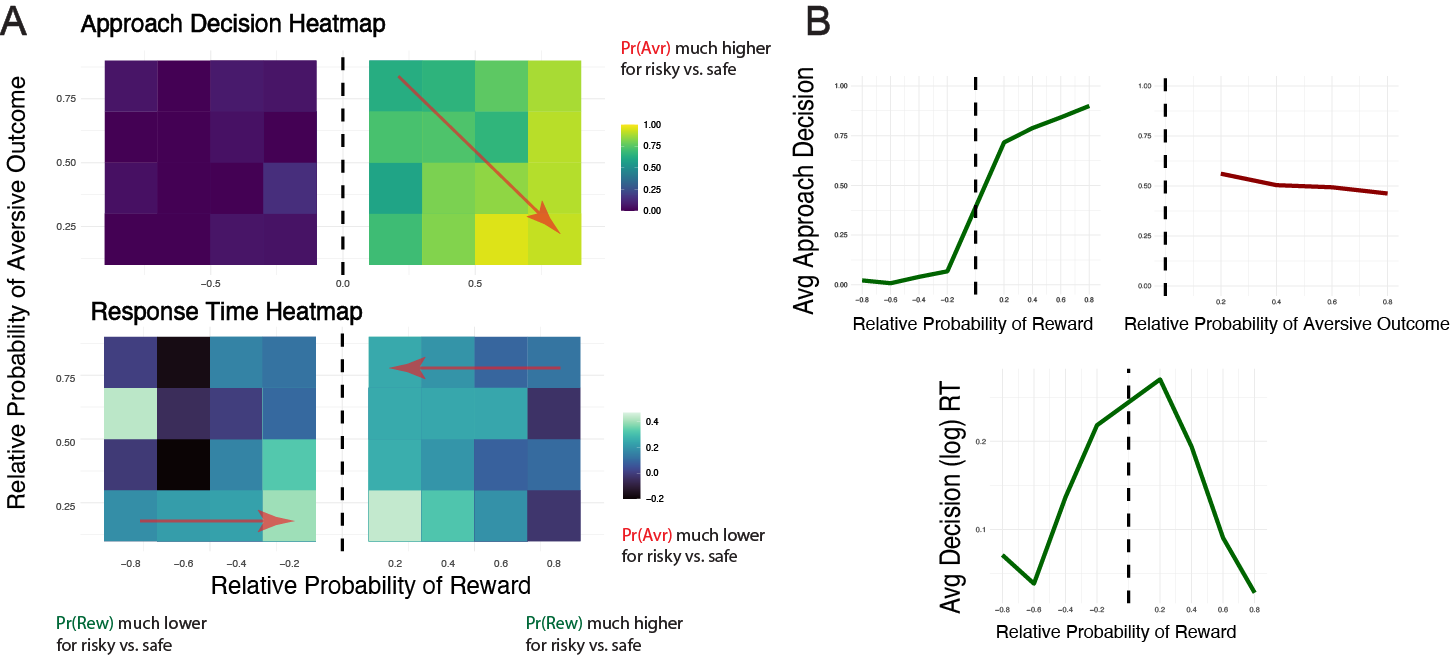

Figure S7. Study 2 behavioral patterns in PAAT.

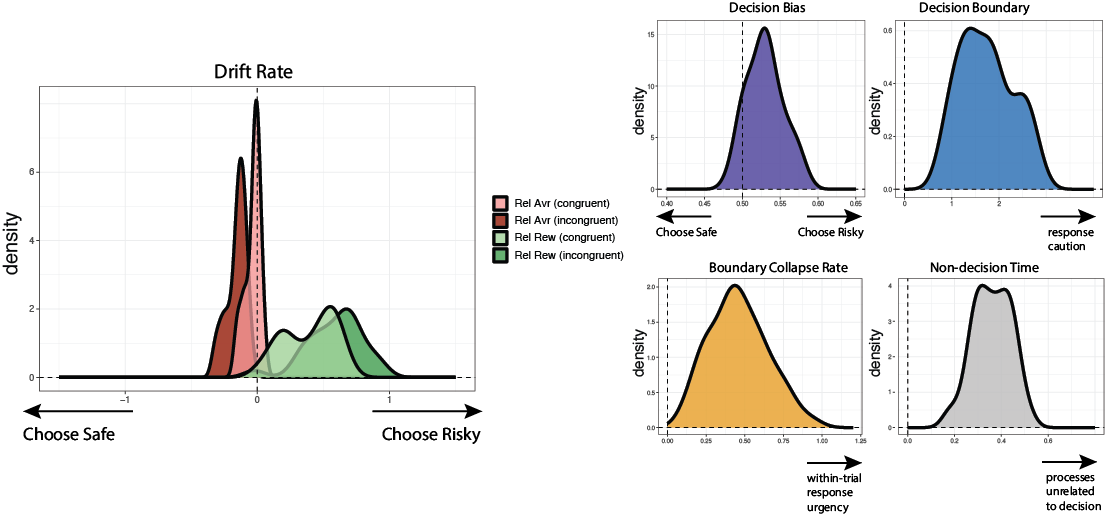

Figure S8. Study 2 group-level posterior distributions of the LCM parameters.

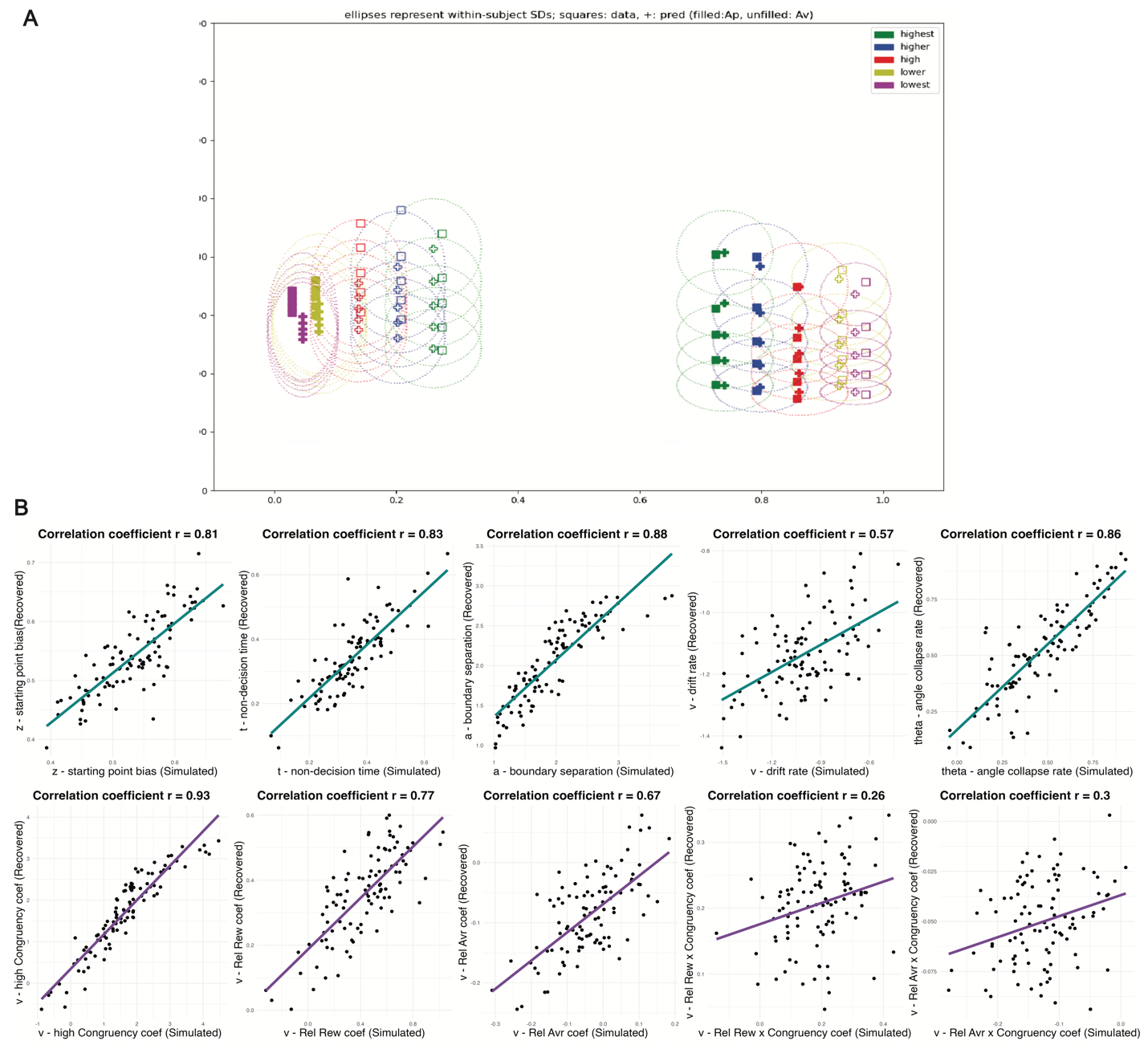

Figure S9. Study 2 Computational Model Validation. (A). Posterior predictive check of the best-fitting model. (B). Simulation results comparing generative parameters (x-axis) with recovered parameters (y-axis).
